# Spatiotemporal dynamics of CD73 in mouse retina under physiological conditions

**DOI:** 10.1101/2025.05.25.655881

**Authors:** Ryutaro Ishii, Keisuke Sakurai, Nao Hosomi, Bernd K. Fleischmann, Seiya Mizuno, Kenichi Kimura, Hiromi Yanagisawa

**Author notes:** Co-Correspondence addressed to: Ryutaro Ishii, M.D., Ph.D. Institute of Medicine, University of Tsukuba. 1-1-1 Tennodai, Tsukuba, Ibaraki 305-8575, Japan, Kenichi Kimura, Ph.D. Life Science Center for Survival Dynamics, Tsukuba Advanced Research Alliance, University of Tsukuba, 1-1-1 Tennodai, Tsukuba, Ibaraki 305-8577, Japan.

## Abstract

Adenosine is essential to energy metabolism and neuromodulation in the central nervous system. The retina is a highly energy-demanding neural tissue, and dysregulation of adenosine signaling causes retinal diseases. However, the dynamics of ecto-5’-nucleotidase (CD73), a key enzyme for extracellular adenosine generation, remain elusive. Here, we investigate its spatiotemporal profile from development to adulthood using two transgenic mouse lines. We found that CD73 is transiently expressed in the early astrocyte lineage (Embryonic day 16.5 to Postnatal day [P] 3), becomes prominent in the rod-photoreceptor lineage by P3, and appears in the inner nuclear layer from P7 onward. CD73 deletion delays rod-response recovery and shortens the implicit time in scotopic ERG under dim light. These findings indicate that light and other physiological cues influence CD73 expression, which has significant consequences for retinal function, thereby providing a foundation for exploring disease-related alterations in CD73 and designing therapies that restore adenosine homeostasis.

## Introduction

Adenosine is essential to cellular energy metabolism as a component of adenosine triphosphate (ATP) and a potent extracellular neuromodulator that shapes neuronal excitability, transmitter release and the sleep–wake cycle (Garcia-Gil et al., 2021; Lazarus et al., 2019; Wei et al., 2011). Extracellular adenosine is produced mainly by ecto-5′-nucleotidase (CD73), which hydrolyzes AMP to adenosine, and is cleared when intracellular adenosine kinase (ADK) re-phosphorylates adenosine imported through equilibrative nucleoside transporters (ENTs). Through these mechanisms, adenosine collectively maintains central nervous system (CNS) homeostasis (Garcia-Gil et al., 2021). Because genetic deletion of adenosine signaling components often triggers compensatory pathways (Garcia-Gil et al., 2021; Wei et al., 2011), dissecting adenosine pathways demands tools with high spatiotemporal and cell-type specificity (Oishi et al., 2017; Wu et al., 2023).

The retina is a high-energy extension of CNS and is continuously exposed to fluctuating levels of light, oxygen, and metabolic substrates (Joyal et al., 2018). Because the ATP is rapidly turned over into ADP and AMP, and ultimately adenosine, metabolic demand directly influences intracellular adenosine dynamics (Garcia-Gil et al., 2021). Moreover, changes in intracellular adenosine during circadian cycle have been shown to govern extracellular adenosine concentrations via ENTs (Cao et al., 2020; Ribelayga & Mangel, 2005). Therefore, in the adult retina, adenosine levels fluctuate in response to daily physiological cues.

As a signalling molecule in the retina, adenosine is well-characterized for its role in pathological angiogenesis such as oxygen-induced retinopathy (OIR) model; adenosine acting via A2A receptors amplifies HIF-1α-dependent angiogenesis by up-regulating glycolysis (Liu et al., 2017). During normal vascular development, however, genetic inactivation of A1 or A2A receptors or CD73 does not cause obvious abnormalities (Liu et al., 2010; Zhang et al., 2022; Zhang et al., 2015). Recent studies have demonstrated that multiple neurotransmitters involved in retinal waves (e.g., dopamine, acetylcholine, and glutamate) (Biswas et al., 2020; Biswas et al., 2024; Liang et al., 2023; Weiner et al., 2019) and light-dependent neural activities influence the retinal vascular system during development (D’Souza & Lang, 2020; Nguyen et al., 2019; Rao et al., 2013). Therefore, adenosine, which modulates stage I and II retinal waves (Huang et al., 2014; Syed et al., 2004; Torborg & Feller, 2005), might contribute to normal retinal vascularization via largely unexplored mechanisms that integrate its intra- and extracellular adenosine metabolism and are driven by physiological neural activity or metabolic cues.

From development through adulthood, intra- and extracellular adenosine metabolism thus play a pivotal role in the retina. Notably, CD73 is abundantly expressed in the rod-photoreceptor lineage (Koso et al., 2009) and modulates extracellular adenosine levels in the OIR model during retinal development and across adult light–dark cycles (Cao et al., 2020; Ribelayga & Mangel, 2005; Zhang et al., 2022). Because the expression patterns of adenosine signalling components and their pathological roles are complex in the retina (Santiago et al., 2020), a detailed characterization of the CD73 profile (its expression and impact on adenosine signalling) under normal conditions is essential for understanding adenosine biology in physiological and pathological contexts. However, the profile remains poorly defined, leaving key aspects of adenosine metabolism unknown.

Here, we mapped the CD73 profile in the retina from embryogenesis through adulthood using newly generated transgenic mouse lines (*CD73-BAC-EGFP* and *CD73-CreER^T2^*). We revealed developmentally regulated patterns of CD73 expression not only in the rod lineage but also in the inner nuclear layer (INL). Moreover, CD73 was transiently expressed in the early astrocyte lineage. CD73 deficiency led to delayed recovery of rod photoresponse and shortened b-wave implicit time under dim-light conditions. The intrinsic fluctuations we uncovered would have remained undetectable by receptor manipulation alone, thereby lay the foundation for investigating how lifestyle factors disturb CD73 function from normal physiology to its pathological states.

## Results

### *CD73-BAC-EGFP* mice reveal that CD73 expression is primarily restricted to the rod lineage, robustly detectable from P3 onward

The retina is dominated by a vast number of rod photoreceptors (Figure 1A) (Swaroop et al., 2010), and previous studies have shown that CD73 is expressed in rods (Koso et al., 2009) and used as a rod-specific marker (Sarin et al., 2018). However, several reports have also noted its presence in other retinal cell types, including Müller glia (Wurm et al., 2011) and the retinal pigment epithelium (RPE) (Chen et al., 2014). To resolve this discrepancy, we first examined CD73 expression in our *CD73-BAC-EGFP* (*CD73-EGFP*) mice (Breitbach et al., 2018) under physiological conditions. In adult retinas, EGFP signals were predominantly localized to the outer nuclear layer (ONL), corresponding to photoreceptors and were undetectable in horizontal cell, bipolar cell, amacrine cell, or Müller glial cells within the INL (Figure 1B). Immunostaining with cone-arrestin (Arr3), a marker of cone photoreceptors (cones), confirmed that EGFP was absent from cones (Figure 1C), indicating that rods are the primary source of EGFP signals in the ONL.

**Figure 1.**
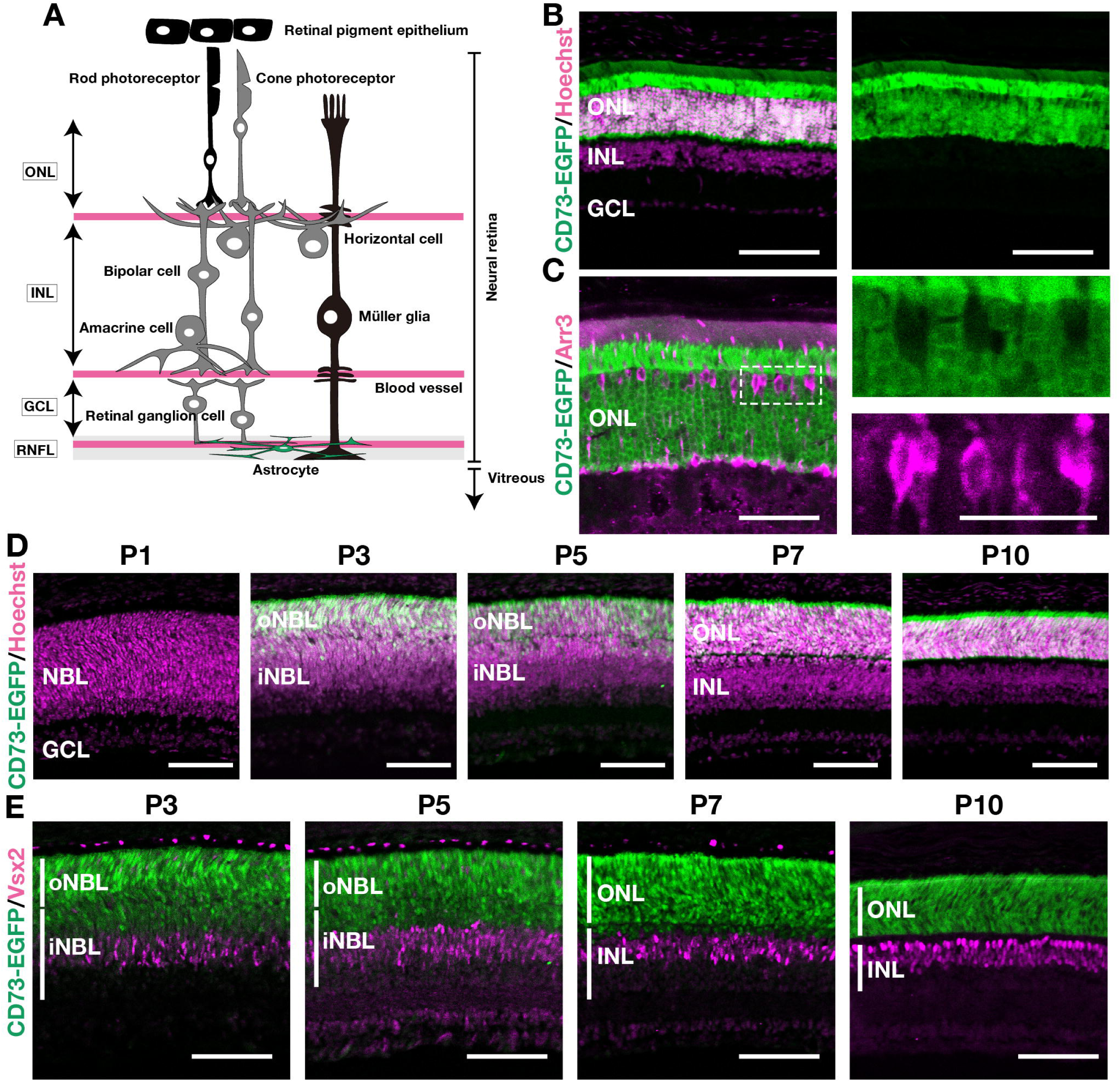
The *CD73-EGFP* mice reveal that CD73 is confined to the rod lineage and becomes detectable from P3 onward. (A) Schematic illustration of adult retinal anatomy. Cell types drawn in black (RPE, Rod photoreceptor and Müller glia) have been previously reported to express CD73. Astrocyte is shown in green and blood vessels in magenta. (B) Representative immunofluorescence images from adult (2–3-month-old) *CD73-BAC-EGFP* mice. Left panel: merged image (CD73-EGFP in green; Hoechst in magenta). Right panel: single-channel image of CD73-EGFP. (C) Higher-magnification views of adult retinas. Left panel: merged image of Arr3 (cone arrestin, magenta) and CD73-EGFP (green), indicating that cones do not express EGFP. Right panel: Higher-magnification view of the dashed box in left panels. The top panels show the CD73-EGFP (green) channel, while the bottom panels show the Arr3 (magenta) channel. (D) Developmental time course (P1–P10) of CD73-EGFP in *CD73-BAC-EGFP* (*CD73-EGFP*) mice retinas. EGFP (green) and Hoechst (magenta) staining illustrate the spatiotemporal emergence of EGFP^+^ cells, notably in the outer neuroblastic layer (oNBL) starting around P3, though some EGFP^+^ cells were observed in the outer part of inner NBL (iNBL). (E) Immunostaining for the bipolar cell marker Vsx2 (magenta) in CD73-EGFP (green) mice retinas. Scale bars: 100 µm (B, D, and E), 50 µm (C left panel), 25 µm (C right panel). Abbreviations: GCL, ganglion cell layer; iNBL, inner neuroblastic layer; INL, inner nuclear layer; NBL, neuroblastic layer; oNBL, outer neuroblastic layer; ONL, outer nuclear layer; RNFL, retinal nerve fiber layer; RPE, retinal pigment epithelium.

We next investigated CD73 expression during development. During early postnatal stages, the neuroblastic layer (NBL) splits into the outer neuroblastic layer (oNBL) and the inner neuroblastic layer (iNBL) around P3; these layers later mature into the ONL and the INL, respectively (Figure 1—figure supplement 1) (Burger et al., 2021). CD73-EGFP expression became robust within the NBL from P3 onward (Figure 1D). Although we observed only minimal EGFP expression in the NBL at P0–P1 in the whole-mount view (Figure 1—figure supplement 2), we could not fully specify the retinal cell types prior to P3. This result was not apparently different from previous reports; CD73 expression increases during the early postnatal days (P0-P4), reaching robust levels by P5 (Koso et al., 2009; Lakowski et al., 2011). Moreover, Arr3 immunostaining confirmed that these EGFP^+^ cells were primarily restricted to the Arr3-negative rod lineage rather than to cone (Figure 1—figure supplement 3). Although EGFP was undetectable in the INL in the adulthood, from P3 to P7, we occasionally detected EGFP signals in both the iNBL and oNBL (Figure 1D). Because the rod lineage originates from the same late progenitor pool as bipolar cells (Brzezinski & Reh, 2015; Wang et al., 2014), we asked whether these transient signals might include bipolar precursors. Immunostaining for Vsx2 (Chx10), a bipolar cell marker, showed that Vsx2-positive cells were EGFP-negative (Figure 1E), indicating that CD73-EGFP is not expressed in the bipolar lineage. These occasional EGFP^+^ cells in the iNBL are therefore most likely rod precursors, consistent with the transient presence of rod lineage cells in this layer during early postnatal development (Burger et al., 2021). Thus, we concluded that our *CD73-EGFP* mice demonstrate that CD73 expression became detectable after P3 in the rod lineage and remains rod-specific in adulthood under physiological conditions.

### *CD73-CreER^T2^;tdTomato* mice further validate that predominant CD73 expression in the neural retina begins in the rod lineage from P3 onward

Next, to confirm and extend our CD73-EGFP observations across multiple developmental stages, we generated another transgenic mouse line by crossing the *CD73-CreER^T2^* knockin mice with Rosa26-loxp-STOP-loxp-tdTomato (tdTomato) knockin mice to obtain *CD73*-*CreER^T2^*; *tdTomato* mice (Figure 2A). We then performed the lineage tracing to determine how *CD73CreER^T2^*^+^ cells at tamoxifen-treated time points differentiate into various cell types at the time of sampling. First, to check for CreER^T2^ leakiness, we examined mice that did not receive tamoxifen and confirmed that no tdTomato signal was detected (Figure 2—figure supplement 1A). Next, to assess the adult expression of *CD73-CreER^T2^*, we administered tamoxifen to 2–3-month-old mice two weeks prior to tissue collection (Figure 2B). In previous studies investigating bone marrow using *CD73-EGFP* mouse, CD73-EGFP expression was observed in the endosteal tissue (Breitbach et al., 2018). Similarly, to validate the accuracy of the Cre mouse model, we examined *CD73-CreER^T2^;tdTomato* mice and found tdTomato-labelled cells in the same endosteal region, consistent with *CD73-EGFP* mice result (Figure 2—figure supplement 1B; arrowheads). Similar to the bone marrow findings, tdTomato-labelled cells in the retina were abundant in the ONL, in agreement with our CD73-EGFP results. Immunostaining for Arr3 confirmed that the tdTomato^+^ cells were Arr3-negative, consistent with rods (Figure 2C). Thus, using two transgenic mouse lines, we confirmed that CD73 is expressed in the rods in the adult retina. Furthermore, in the *CD73*-*CreER^T2^*; *tdTomato* mice, we also observed a small population of tdTomato^+^ cells in RPE65+ RPE (Figure 2—figure supplement 1C), which was not clearly detected in the *CD73-EGFP* mice. This discrepancy may reflect the stronger tdTomato signal in *CD73-CreER^T2^;tdTomato* mice compared with the EGFP signal in *CD73-EGFP* mice (Figure 2—figure supplement 1B), which enables more sensitive detection of CD73^+^ cells.

**Figure 2.**
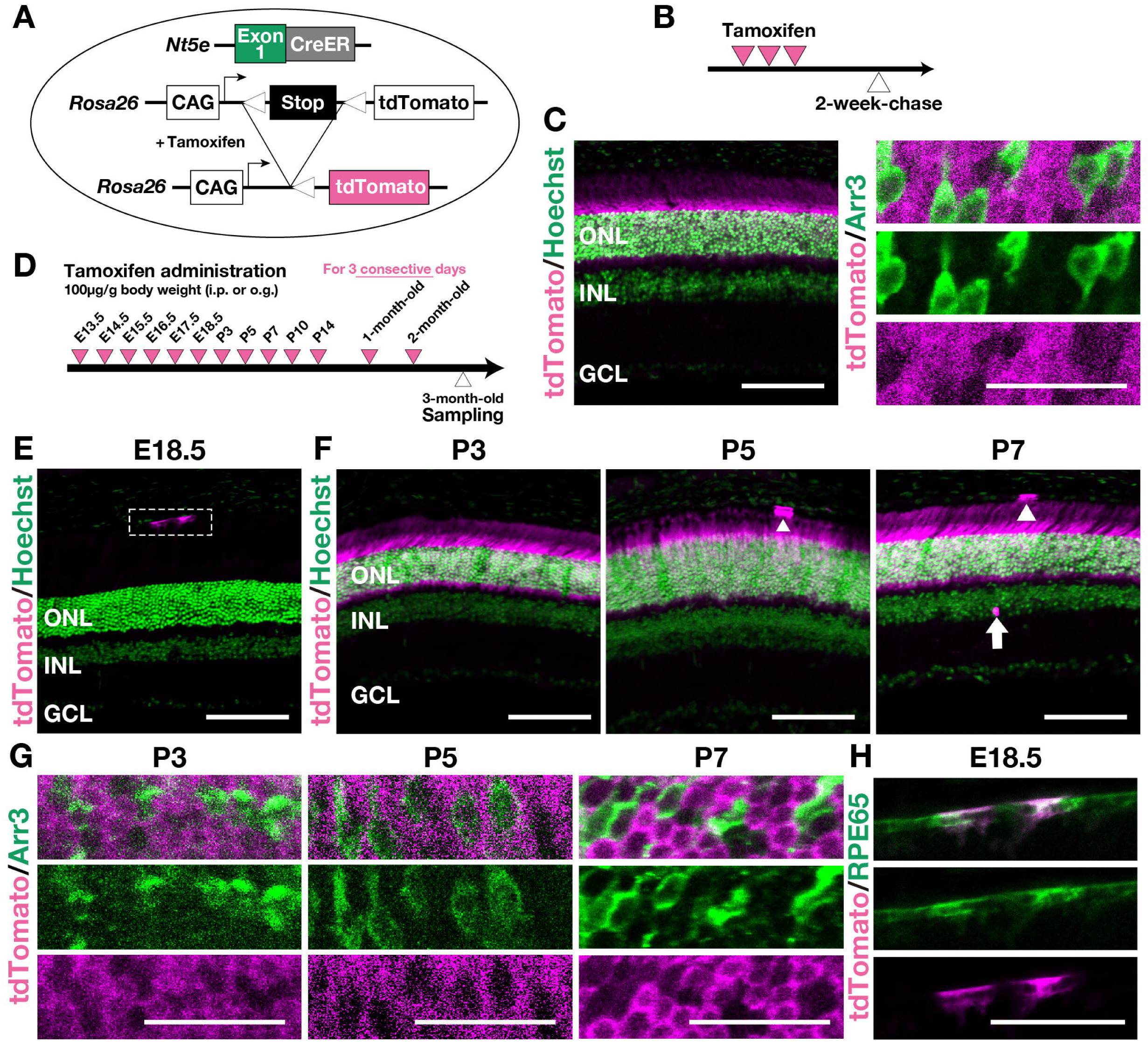
Lineage tracing experiments confirm that CD73 expression in the ONL is restricted to the rod lineage from P3 onward. (A) Schematic of the tamoxifen-inducible *CreER^T2^* system for labeling CD73-fated cells. (B) Experimental timeline showing tamoxifen injections (magenta arrowheads) and sample collection (white arrowheads) in adult mice. (C) Representative images of adult *CD73-CreER^T2^;tdTomato* retinas two weeks after tamoxifen injection. Left panel: Hoechst (green) labels nuclei. Right panel: higher-magnification view of the outer portion of the ONL. Cone photoreceptors are labeled with Arr3 (cone arrestin, green). The top row shows the merged image, the middle row shows Arr3 alone, and the bottom row shows tdTomato alone. tdTomato is shown in magenta. (D) Schematic of the lineage-tracing strategy to visualize descendants of *CD73-CreER^T2+^*cells from E13.5 to 2 months of age, with tissue collection at 3 months of age. Magenta arrowheads indicate tamoxifen injection days for each of the experimental groups; white arrowheads mark tissue collection. Further details are provided in the Materials and Methods. (E, F, G, H) Lineage tracing in *CD73-CreER^T2^;tdTomato* mice treated with tamoxifen at E18.5, P3, P5, or P7 and analyzed at 3 months of age. Panels (G) and (H) show higher-magnification views of (F) and (E), respectively. In (E), the white dashed boxes outline the regions shown in (H). In (E) and (F), Hoechst (green) labels nuclei; in (H), RPE65 (green) marks the RPE; in (G), Arr3 (green) marks cone photoreceptors. tdTomato is shown in magenta. tdTomato^+^ cells are absent in the neural retina (ONL, INL, and GCL) at E18.5 (E), but they are predominantly located in the ONL at P3, P5, and P7 (F). The dashed box in (E) shows tdTomato^+^ cells in the RPE (E), illustrated at higher magnification in (H). Abbreviations: GCL, ganglion cell layer; INL, inner nuclear layer; ONL, outer nuclear layer; RPE, retinal pigment epithelium. Scale bars: 100 µm (C, left; E; F); 50 µm (H); 25 µm (C, right; G)

We then investigated earlier developmental stages by performing lineage tracing experiments. We administered tamoxifen at various time points from E13.5 to 2 months of age and examined the retinas at 3 months of age (Figure 2D). When tamoxifen was given at E18.5, the tdTomato signal was not apparent in the ONL (Figure 2E). However, tdTomato^+^ cells were clearly visible in the ONL when tamoxifen was administered after P3 (Figure 2F). In contrast, few tdTomato^+^ cells appeared in the INL in any tamoxifen-treated group (Figure 2F; Figure 2—figure supplement 2A). Furthermore, we confirmed that these tdTomato^+^ cells in the ONL were Arr3-negative (Figure 2G; Figure 2—figure supplement 2B), indicating that the *CD73-CreER^T2^* is expressed in the rod lineage. Although we could not fully characterize their distribution from transverse sections alone, we did detect tdTomato^+^ RPE in tamoxifen-treated groups from embryonic stages onward (Figures 2E, 2F and 2H, Figure 2—figure supplement 1C, 2A and 2C; arrowheads), suggesting that CD73 expression in the RPE is regulated by mechanisms distinct from those operating in the neural retina. Collectively, these lineage-tracing data reinforce our CD73-EGFP findings, demonstrating that CD73 expression in the neural retina is readily detectable in the rod lineage by P3 and persists into adulthood.

### CD73 is transiently expressed in the early astrocyte lineage from E16.5 to P3, and becomes apparent in the INL at P7, persisting into adulthood

In transverse sections, a small number of tdTomato^+^ cells were also observed in the INL (Figure 2F, Figure 2—figure supplement 2A; arrow) and in the retinal nerve fiber layer (RNFL), the thin layer immediately vitreal (inner) to the GCL (Figure 2—figure supplement 2D; arrowheads; see Figure 1A for an anatomical schematic). To further clarify the spatiotemporal distribution of these tdTomato^+^ cells in the INL and RNFL, we examined whole-mount retinas from *CD73-CreER^T2^;tdTomato* mice. We found that tdTomato^+^/S100β^+^ (an astrocyte marker) cells in the RNFL were present during embryonic and early postnatal stages (Figure 3A), whereas a few tdTomato^+^ cells in older retinas appeared in the INL (Figure 3B). Most tdTomato^+^ somata in the INL were round, and lay on the side facing the GCL (Figure 3B, arrowheads); thereby, based on their position and morphology (Duda et al., 2025; Yan et al., 2020), we consider them amacrine cells. However, a few tdTomato^+^ cells exhibited distinct shapes (Figure 3B, right panel, arrow). Three-dimensional reconstructions confirmed that these cells were Müller glia spanning the ONL to the GCL (Figure 3C, arrow). These results indicated CD73^+^ cells in the INL were mainly amacrine cells and Müller glia, although the small number of CD73^+^ cells precluded a more precise assessment of their cell-type specificity.

**Figure 3.**
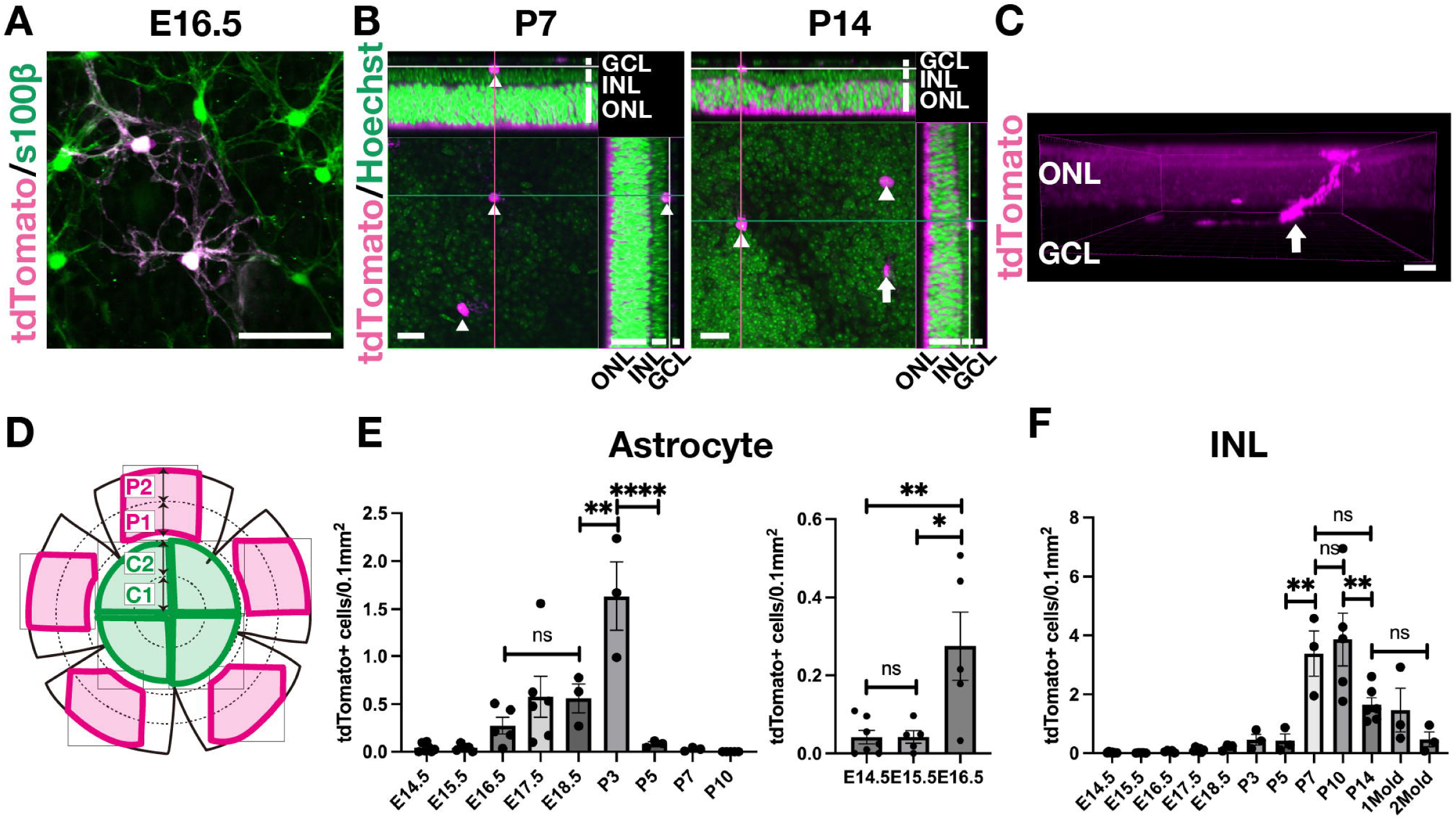
Temporal distribution of descendants of *CD73-CreER^T2+^* cells in the RNFL and INL at 3 months of age. (A) Representative image of tdTomato^+^ astrocytes in the RNFL. Tamoxifen was administered at E16.5. S100β (an astrocyte marker) is labeled in green and tdTomato is magenta. (B) Orthogonal views of representative tdTomato^+^ cells in the INL. Tamoxifen was administered at P7 (left) or P14 (right). The main panel shows the xy plane; the xz (top) and yz (right) planes are orthogonal slices (green and magenta lines, respectively; white lines in xz and yz indicate the z-position of the xy plane). Hoechst (green) labels nuclei, and tdTomato is shown in magenta. Arrowheads indicate tdTomato^+^ cells located the GCL-facing side the INL, consistent with the typical distribution of amacrine cells. (C) 3D Imaris reconstructions of tdTomato^+^ cells at P14. tdTomato is shown in magenta. The tdTomato^+^ cell indicated by the arrow corresponds to the one marked by the arrow in (B, right panel). (D) Schematic illustrating tdTomato^+^ cell quantification. Whole-mount retinas were divided into central (green) and peripheral (magenta) regions. The central region was split into C1 (0–500 μm from the optic nerve head) and C2 (500–1000 μm), while the peripheral region was split into P1 (500–1000 μm from the periphery) and P2 (0–500 μm). tdTomato^+^ cells were counted in each zone. Additional details are provided in the Materials and Methods. (E, F) Temporal characteristics of tdTomato^+^ cells in astrocytes (E) and the INL (F) in different tamoxifen-treatment groups (E14.5 to P10 in E, left panel; E14.5 to E16.5 in E, right panel; E14.5 to 2 months of age in F). Data are presented as mean ± s.e.m., with each dot representing an individual sample (n ≥ 3 retinas per condition, one retina per mouse). One-way ANOVA was used to compare tdTomato^+^ cell densities among time points, followed by Tukey’s multiple comparisons. All statistical analyses were performed in GraphPad Prism v10 (GraphPad Software). Significance: * *p* < 0.05; ** *p* < 0.01. Detailed calculation methods are provided in the Material and Methods. Abbreviations: GCL, ganglion cell layer; INL, inner nuclear layer; ONL, outer nuclear layer; RNFL, retinal nerve fiber layer. Scale bars: 50 μm (A; B), 20 μm (C).

To investigate these patterns in the astrocyte lineage and the INL, we performed quantitative analyses of the entire retina from the optic nerve head to the periphery (Figure 3D; see Figure 3—figure supplement 1 for anatomical details). First, we examined the temporal changes in descendants of CD73^+^ cells in the astrocyte (tdTomato^+^/S100β^+^ cells). The number of tdTomato^+^ cells increased until P3, then sharply declined by P5 and P7; by P10 no cells were detected (Figure 3E, left panel). Although the density of tdTomato^+^ cells differed markedly between E18.5 and P3, tamoxifen administration differed between these stages (oral gavage to the pregnant dam at E18.5 versus to the pups at P3), making a direct comparison unreliable. Although a few tdTomato^+^ cells were present at E14.5 and E15.5, their density increased substantially from E16.5 onward (Figure 3E, right panel). By contrast, the number of lineage-traced descendants of CD73^+^ cells (tdTomato^+^ cells) in the INL increased gradually until P5, then surged at P7, and persisted into adulthood with only a slight decline thereafter (Figure 3F).

We next investigated the spatial distribution of tdTomato^+^ (descendants of *CD73-CreER^T2+^*) astrocytes in 3-month-old *CD73-CreER^T2^;tdTomato* mice. Tamoxifen was administered at E16.5, E17.5, E18.5, or P3, which correspond to the peak density of tdTomato^+^ astrocytes (Figure 3E). When tamoxifen was given at E16.5, descendants of *CD73-CreER^T2+^* cells migrated centrifugally to the periphery (P1 and P2; Figures 4A and 4B) rather than remaining in the central regions (C1 and C2; Figure 4—figure supplement 1), although no statistically significant differences were found among these regions (Figure 4C). In contrast, tdTomato^+^ descendants were detected at statistically significant levels in the most peripheral areas for mice treated at E17.5, E18.5, or at P3 (Figure 4 and Figure 4—figure supplement 1). Finally, we examined tdTomato^+^ cells in the INL starting at P7, when their number began to increase (Figure 3F). Although the tdTomato^+^ cells in the INL appeared slightly more concentrated in the central region (Figure 5A, 5B; Figure 5—figure supplement 1), there were no significant differences among all regions (Figure 5C).

**Figure 4.**
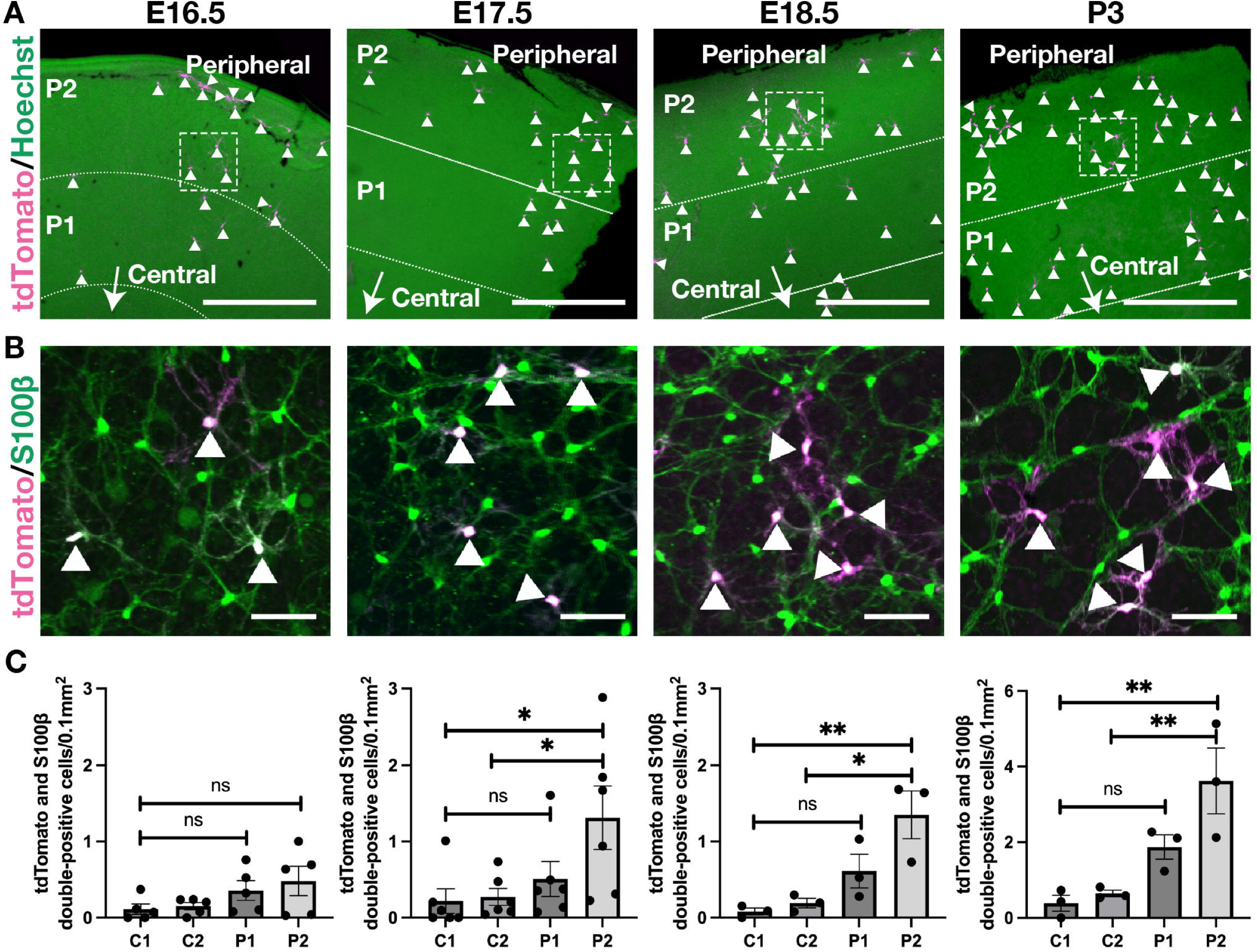
Descendants of *CD73-CreER^T2+^* cells in the astrocyte lineage are enriched in the peripheral retina. (A) Representative whole-mount retinal images from *CD73-CreER^T2^;tdTomato* mice treated with tamoxifen at E16.5, E17.5, E18.5, or P3. Maximum-intensity projection images spanning from the INL to the RNFL, excluding the ONL, provide overviews of the peripheral retina for each tamoxifen treatment group. Hoechst (green) labels nuclei, and tdTomato is shown in magenta. “Peripheral” marks the retinal periphery, and the arrow labeled “Central” indicates the direction toward the optic nerve head. (B) Higher-magnification views of the white dashed boxes in (A). S100β (an astrocyte marker) is labeled in green and tdTomato is magenta. Arrowheads indicate tdTomato and S100β double-positive cells. (C) The spatial distribution profiles of tdTomato^+^ astrocytes, measured from the optic nerve head (0 µm) to the peripheral retina (covering C1, C2, P1, and P2). Data are shown as mean ± s.e.m., with each dot representing an individual sample (n = 3 for E16.5, n = 6 for E17.5, n = 3 for E18.5, n = 3 for P3). One-way ANOVA was used to compare the density of tdTomato and S100β double-positive astrocytes across these regions, followed by Tukey’s multiple comparisons. All statistical analyses were performed in GraphPad Prism v10 (GraphPad Software). Significance: * *p* < 0.05; ** *p* < 0.01. Detailed calculation methods are provided in the Materials and methods. Abbreviations: INL, inner nuclear layer; ONL, outer nuclear layer; RNFL, retinal nerve fiber layer. Scale bars: 500 µm (A); 50 µm (B).

**Figure 5.**
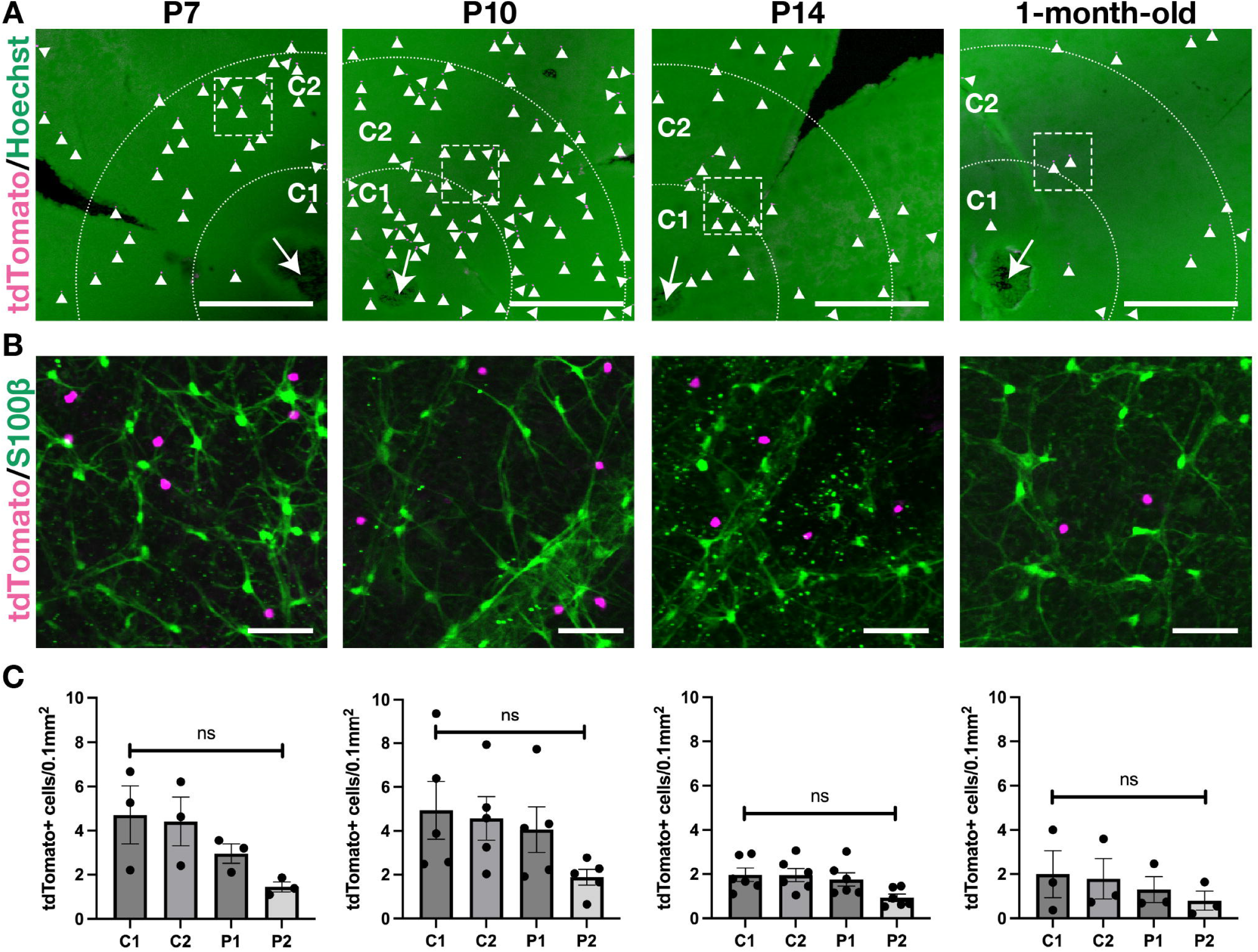
Spatial distribution patterns of descendants of *CD73-CreER^T2+^* cells in the INL. (A) Representative wholemount retinal images from *CD73-CreER^T2^;tdTomato* mice treated with tamoxifen at P7, P10, P14, or 1 month of age. Shown are maximum-intensity projection images from the INL to the RNFL, excluding the ONL, providing an overview of the central retina for each treatment group. Nuclei are labeled with Hoechst (green), and tdTomato is shown in magenta. Arrowheads indicate tdTomato^+^ cells in the INL. (B) Higher-magnification views of the white dashed boxes in (A). The astrocyte marker S100β is labeled in green, and tdTomato is shown in magenta. These images confirm that the tdTomato^+^ cells in (A) reside in the INL and lack S100β expression. (C) The density of tdTomato^+^ cells in the INL, measured from the optic nerve head (0 µm) to the peripheral retina. Data are shown as mean ± s.e.m., with each dot representing an individual sample (n = 3 for P7, n = 5 for P10, n = 6 for P14, n = 3 for 1-month-old). One-way ANOVA was used to compare the density of tdTomato^+^ cells in the INL across the four regions (C1, C2, P1, P2), followed by Tukey’s multiple comparisons. All statistical analyses were performed in GraphPad Prism v10 (GraphPad Software). Detailed calculation methods are provided in the Materials and Methods. Abbreviations: INL, inner nuclear layer; ONL, outer nuclear layer; RNFL, retinal nerve fiber layer. Scale bars: 500 µm (A); 50 µm (B).

Taken together, our lineage-tracing data indicate that CD73 is expressed within the early astrocyte lineage and its descendants accumulate in the periphery, whereas CD73^+^ cells in the INL are distributed throughout the retina with a slight central enrichment.

### CD73 is dispensable for retinal cell genesis under physiological conditions

A previous in vitro study suggested that CD73 is not required for rod-lineage differentiation (Koso et al., 2009). However, recent studies in the retinal vasculature and in other tissues have shown that receptor-independent adenosine signalling that drives epigenetic modifications (Garcia-Gil et al., 2021; Xu et al., 2017). Therefore, we generated *CD73-CreER^T2^* knock-in/knock-out (CD73 KO) mice by mating *CD73-CreER^T2^* mice to homozygosity (Figure 5—figure supplement 2) and examined retinal cell genesis by quantifying transverse sections at P14, as described previously (Zhang et al., 2023). Our analyses with *CD73-BAC-EGFP* and *CD73-CreER^T2^;tdTomato* mice showed that CD73 becomes distinctly detectable in the rod lineage by P3 (Figure 1D, E; Figure 1—figure supplement 3; Figure 2F, G; Figure 2—figure supplement 2A-C), suggesting that late-born retinal cells (bipolar cells, Müller glia, and rods) might be influenced by CD73 activity. Because most tdTomato^+^ somata in the INL were considered to be amacrine cells (Figure 3B), we also confirmed their identity. Immunostaining for Vsx2^+^ bipolar cells, Lhx2^+^ Müller glia, and AP2α^+^ amacrine cells (Figure 5—figure supplement 3A) revealed no statistically significant differences among these cell types between wild-type (WT) and CD73 KO retinas (Figure 5—figure supplement 3B). Moreover, we performed H&E staining to evaluate the rod development by measuring the ONL thickness, which was comparable across genotypes (Figure 5—figure supplement 3C). These findings indicate that, under physiological conditions, CD73 deficiency does not affect retinal cell genesis.

### The impact of CD73 dysfunction on retinal photoresponses is prominent under scotopic conditions but becomes negligible under photopic conditions

In the adult retina, most extracellular adenosine under physiological conditions is generated via the CD73 pathway, and these levels change during light/dark adaptation (Ribelayga & Mangel, 2005). A recent study showed that intravitreal administration of a CD73 inhibitor in dark-adapted C57BL/6N mouse eyes had no effect on a-wave or b-wave amplitudes (Losenkova et al., 2022). However, whether the complete loss of adenosine production by CD73 induces physiological and morphological changes in the mouse retina under normal light conditions remains unclear. To investigate the impact of CD73-mediated adenosine on retinal function, we performed scotopic and photopic ERG analyses in WT and CD73 KO mice. In the scotopic ERG, the average waveforms evoked by various flash intensities are shown (Figure 6A). At moderate or bright light intensities, photoresponses were comparable between WT and CD73 KO mice (Figure 6A, middle and right). However, under the dim light conditions, CD73 KO mice exhibited slightly accelerated response kinetics relative to WT mice (Figure 6A, left). Notably, the amplitudes of the a-wave and b-wave, which primarily reflect rod photoreceptors and ON bipolar cells, respectively, were not significantly different between the two genotypes (Figure 6B, left and middle). In contrast, the implicit time (time-to-peak) of the b-wave, particularly under dim light conditions, was modestly shorter in CD73 KO mice compared to WT mice (Figure 6B, right). Additionally, the initial activation phase of the a-wave was virtually identical between WT and CD73 KO mice, suggesting no effect on the activation processes of the rod phototransduction cascade (Figure 6—figure supplement 1). In the photopic ERG, where the rod responses are suppressed by background illumination and the waveforms primarily reflect cone function, the amplitudes and implicit times of photopic b-waves were similar between CD73 KO and WT mice (Figure 6C, D). These findings indicate that CD73-derived adenosine activity, while negligible under bright light conditions, plays a more pronounced role under low-light conditions. Collectively, in vivo ERG analysis suggests that the adenosine produced by CD73, which is expressed in rod photoreceptors, as confirmed by the results of our *CD73-EGFP* and *CD73-CreER^T2^;tdTomato* mice (Figures 1C and 2C), contributes more prominently to rod-driven retinal responses than to cone-driven ones.

**Figure 6.**
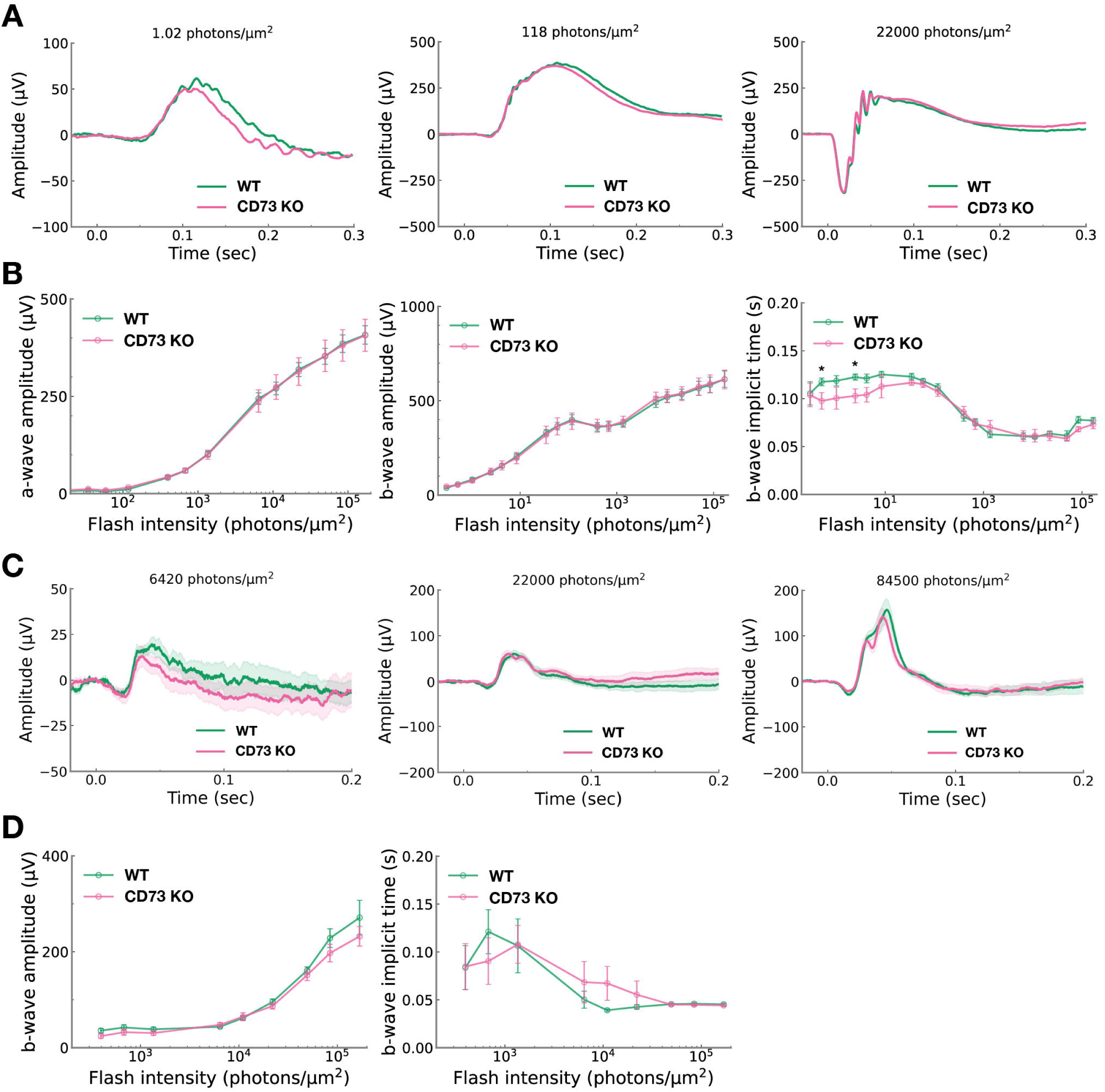
Scotopic and photopic ERG recordings in WT and CD73 KO mice. (A) The average photoresponses recorded from WT (green line, n =10) and CD73 KO mice (magenta line, n = 10) under scotopic conditions. The flash stimuli at time 0 were delivered with photon densities of 1.02 (left), 118 (middle), or 22,000 (right) photons/μm^2^ on the cornea surface. (B) The plot of a-wave amplitude (left), b-wave amplitude (middle), and b-wave implicit time (right) in the scotopic ERG as a function of light intensity for WT (green, n = 10) and CD73 KO (magenta, n = 10) mice. Data are presented as mean ± s.e.m. (C) The average photoresponses recorded from WT (green line, n =9) and CD73 KO mice (magenta line, n = 10) under photopic conditions. The intensity of flash stimuli on the cornea surface is 6,420 (left), 22,000 (middle), and 84,500 (right) photons/μm^2^. The shaded band around each mean response trace represents the ± s.e.m. (D) The plot of b-wave amplitude (left) and b-wave implicit time (right) in the photopic ERG as a function of light intensity for WT (green, n = 9) and CD 73KO (magenta, n = 10) mice. Data are shown as mean ± s.e.m. Statistical significance was assessed using unpaired two-tailed Student’s *t*-test (\**p* < 0.05). Abbreviations: KO, knockout; WT, wild-type.

### CD73 modulates rod recovery under low-light conditions

To directly examine the influence of CD73 on the physiological function of rod photoreceptors, we performed single-cell recordings from individual rods of CD73 KO and WT mice (Figure 7A). The maximal response amplitude (R_max_) and half-saturating response intensities (I_1/2_), which indicates the sensitivity of rod, were comparable between WT and CD73 KO rods (Figure 7B-E). Under dim-light conditions, while the time-to-peak of the dim flash responses was unaffected (Figure 7F), the integration time was significantly prolonged in CD73 KO rods compared with WT rods (Figure 7G). This extended integration time may explain the shorter implicit time of b-wave observed in ERG recordings under dim light conditions. Since the ERG a-wave, which originates from photoreceptors, is a negative-going response, its delayed recovery may prolong the suppressive influence on the subsequent positive-going b-wave after its peak. This could result in an apparently earlier peak of the b-wave, even if the bipolar kinetics are unchanged. The prolonged integration time observed in CD73 KO rods may reflect a delay in the recovery processes of the rod phototransduction cascade, potentially due to rhodopsin phosphorylation by GRK1 or reduced the GTPase activity of the transducin-PDE complex.

**Figure 7.**
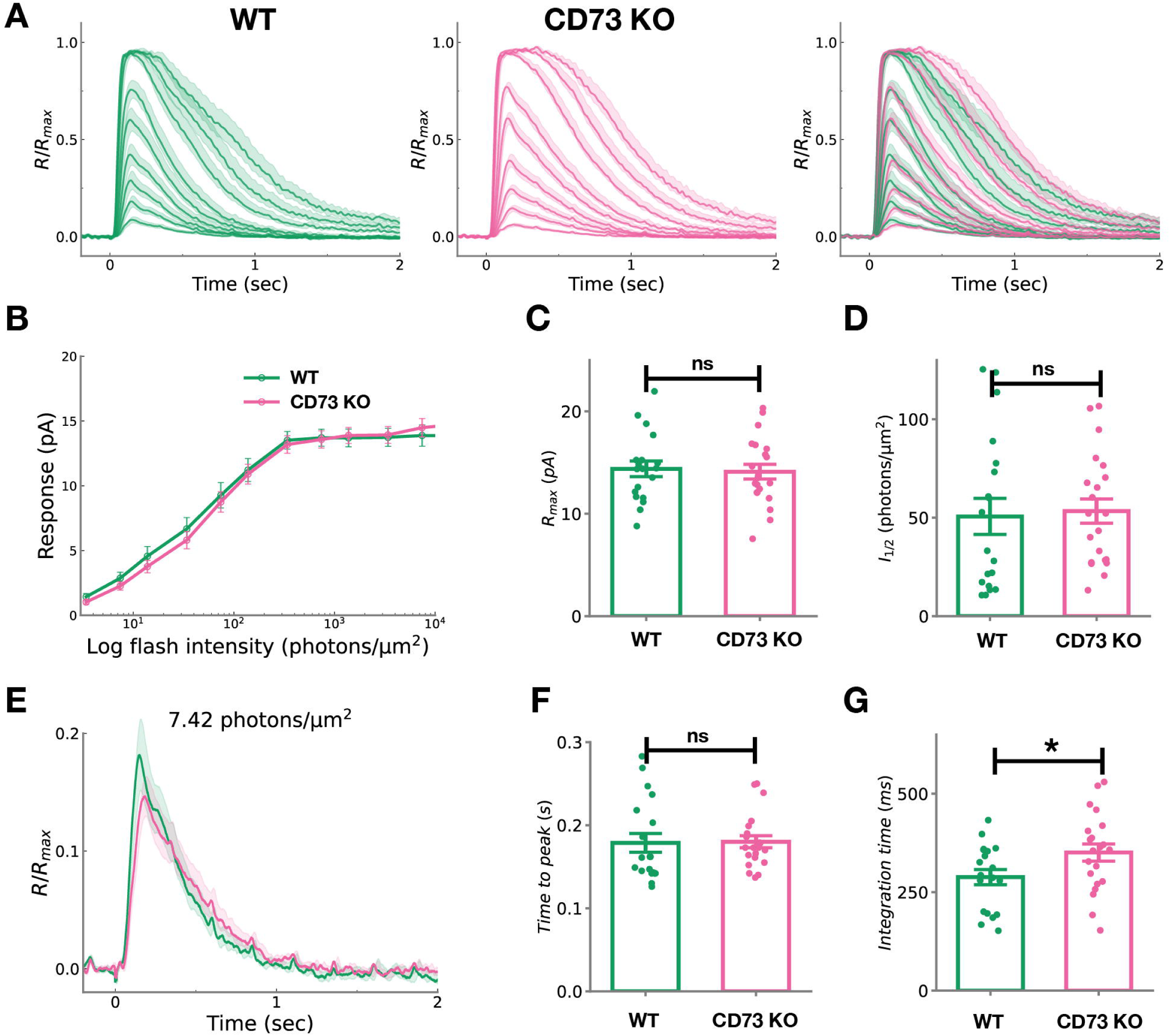
Single-cell recordings of WT and CD73-KO rod photoreceptors. (A) Average response families of WT (left, n = 19) and CD73 KO (middle, n = 21) rod photoreceptors to flashes of varying intensities: 3.4, 7.42, 13.8, 34, 74.2, 138, 340, 742, 1380, and 3400 photons/μm^2^. The right panel shows the merged responses of WT and CD73 KO rods. Responses from each rod photoreceptor were normalized to the maximal response amplitude of the respective photoreceptor. The shaded band around each mean response trace represents the ± s.e.m. (B-D) The intensity-response relationship in WT (green, n = 19) and CD73 KO (magenta, n = 21) rod photoreceptors (B). The maximal amplitude (C) of the response and I_1/2_ (D) in WT and CD73 KO rod photoreceptors. Data are shown as mean ± s.e.m. (E-G) Average response evoked by a flash (7.42 photons/μm^2^) of WT (green, n = 19) and CD73 KO (magenta, n = 21) rod photoreceptors (E). The time-to-peak (F) of the response and integration time (G) of the dim-flash response in WT and CD73 KO. The shaded band around each mean response trace represents the ± s.e.m. (E). Data are shown as mean ± s.e.m., with each dot representing an individual sample (F and G). Statistical significance was assessed using unpaired two-tailed Student’s *t*-test (\**p* < 0.05). Abbreviations: KO, knockout; WT, wild-type.

Next, at P14 and 1 month of age, overall retinal morphology did not differ between WT and CD73 KO (Figure 5—figure supplement 3C; dashed line in Figure 7—figure supplement 1A). At the region located 1000 µm inferior to the optic nerve, ONL thickness at 1 month of age was comparable between WT (56.8 ± 2.3 µm, n = 11) and CD73 KO (55.6 ± 2.9 µm, n = 7) mice (unpaired two-tailed Student’s *t*-test, *p* = 0.75). By 3 months, however, the ONL was significantly thicker in CD73 KO mice (49.0 ± 1.3 µm, n = 9) than in WT (43.4 ± 1.3 µm, n = 10) (unpaired two-tailed Student’s *t*-test, *p* = 0.008; solid line in Figure 7—figure supplement 1A). Although this inferior region is known to receive the highest light exposure and shows light-induced degeneration in *Crb1* mutations (e.g., *Crb1^-/-^* and *Crb1^rd8/rd8^*) (Alves et al., 2014; Mehalow et al., 2003; van de Pavert et al., 2004; van de Pavert et al., 2007), we did not observe comparable ONL disorganization in CD73 KO mice (Figure 7—figure supplement 1B).

Taken together, our results indicate that CD73 deficiency led to delayed recovery of rod photoresponse and affects total ONL thickness in the highest light exposure region.

## Discussion

In this study, we identified two key characteristics of the CD73 profile by employing two novel transgenic mouse lines. First, we delineated the spatiotemporal pattern of CD73 expression from embryogenesis through adulthood, demonstrating that CD73, in addition to its well-established presence in rods, is also expressed in the INL and transiently in the early astrocyte lineage. Second, we detected a diminished rod photoresponse following CD73 disruption (Figure 8). These two findings imply that various factors may regulate CD73 profile under physiological conditions.

**Figure 8.**
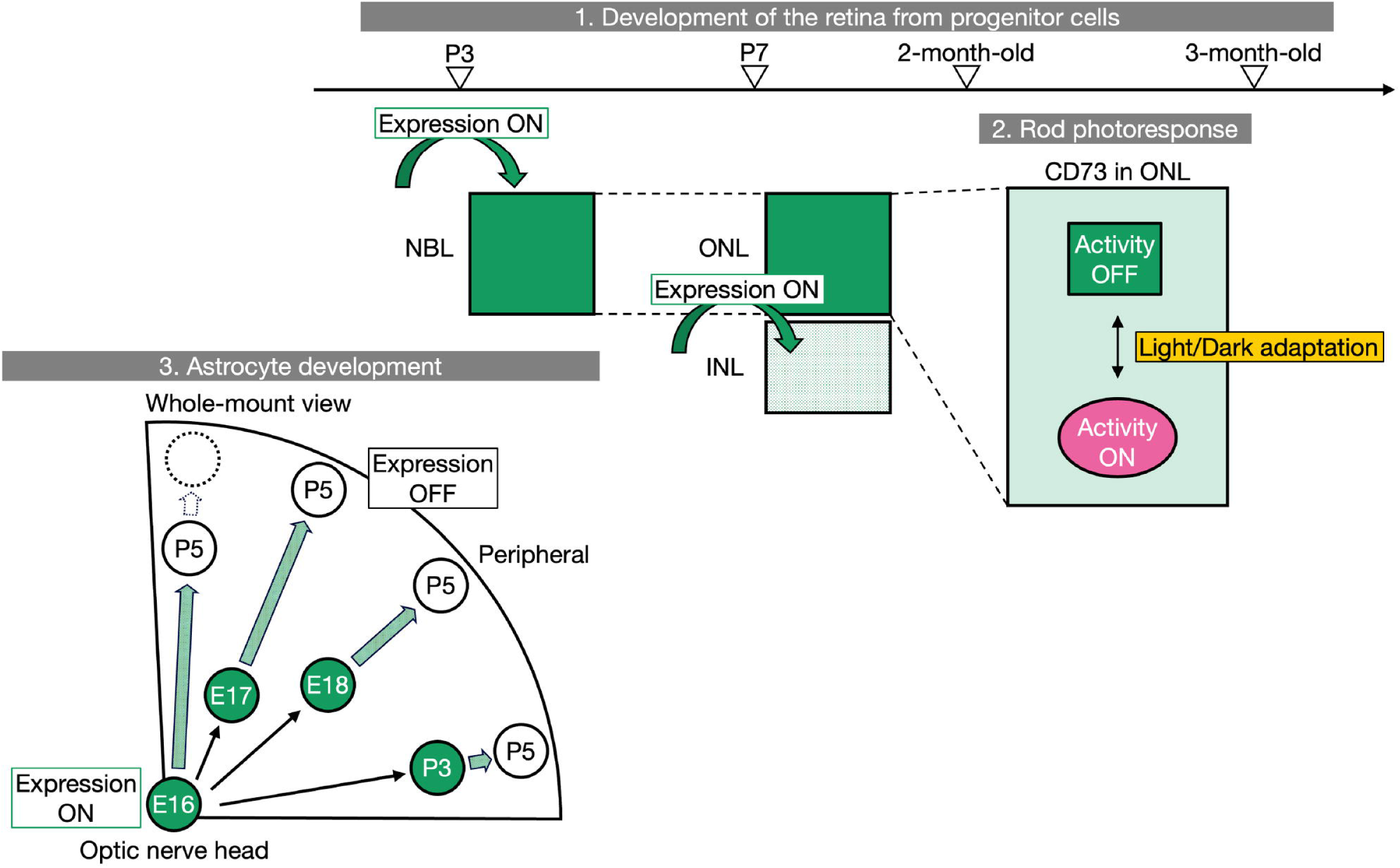
Proposed model of CD73 dynamics in the mouse neural retina under physiological conditions. Spatiotemporal CD73 dynamics follows three patterns. (1) From P3 onward it is abundant in the rod lineage; by P7 it is also initiated in a subpopulation of cells in the INL, peaking at P10 and gradually declining thereafter. CD73 expression in both rods and the INL persists into adulthood. (2) CD73 activity in rods becomes detectable (“ON”) under dim-light conditions, whereas it remains undetectable (“OFF”) under brighter conditions. (3) From E16.5 to P3, CD73 is expressed in a subpopulation of the astrocyte lineage that migrate all the way to the periphery, although its expression is rapidly lost after P3. CD73^+^ cells initially appear around E16.5 in the optic nerve head; however, their descendants do not statistically significantly accumulate in the adult retinal periphery. Abbreviations: INL, inner nuclear layer; NBL, neuroblastic layer; ONL, outer nuclear layer.

### Hypoxia as an upstream regulator of the CD73 profile in the retina

CD73 transcription is directly induced by HIF-1α in various tissues (Alcedo et al., 2021; Synnestvedt et al., 2002). Neonatal retinal HIF-1α mRNA levels rise during the early-postnatal window (Caprara et al., 2011), paralleling the onset of our CD73 expression in the rod lineage from P3 onward. Therefore, HIF-1α is a plausible driver of CD73 expression. However, immunohistochemical mapping of stabilized HIF-1α protein has so far been documented mainly in Pax6^+^ retinal progenitor cells of the inner retina (at P6) (Nakamura-Ishizu et al., 2012); whether HIF-1α similarly accumulates in the rod lineage remains unclear. Furthermore, our adult functional data show that rod photoreceptors, which consume substantial oxygen in darkness (Joyal et al., 2018), rely on CD73 for proper photoresponses. Taken together, these findings suggest that oxygen-dependent adenosine metabolism, especially via the HIF-1α pathway, shapes the CD73 profile from development through adulthood and warrant deeper investigation.

### Light-dependent regulation of adenosine metabolism implied by the temporal CD73 expression during development

Adenosine modulates spontaneous Stage I (E16.5–P0/1) and Stage II (P0/1–P10) retinal waves (Huang et al., 2014; Syed et al., 2004; Torborg & Feller, 2005). The onset of CD73 in the astrocyte lineage at E16.5 coincides with the Stage I initiation, suggesting extracellular CD73-generated adenosine could modulate these waves. By contrast, the later time points (P3, when CD73 expression in the astrocyte lineage declines, and P7, when it begins in the INL) do not align with either the onset or offset of retinal waves, implying that CD73 is regulated by mechanisms independently of retinal-wave activity during postnatal development.

Recent studies have shown that the vitreous hyaloid vasculature undergoes light-dependent remodeling, mediated first by the OPN4–VEGFA pathway (E16–P3) and later by the OPN5–dopamine pathway (P3–P8); OPN4 and OPN5 are expressed in distinct populations of RGC (D’Souza & Lang, 2020; Nguyen et al., 2019; Rao et al., 2013). The pathway switch at P3 coincides with the loss of CD73 in the astrocyte lineage. Because dopamine and adenosine generally work in opposite directions (Cameron et al., 2020; Li et al., 2013), the loss of CD73 at P3 could be significant for the onset of dopamine release by dopaminergic amacrine cells. In addition, the initiation of phototransduction in photoreceptors around P8 (Bonezzi et al., 2018; Tiriac et al., 2018) coincides with the onset of CD73 expression in the INL at P7. Collectively, these observations point to light stimulation as an upstream cue that regulates CD73 expression during development. A recently described light-dependent, retrograde excitatory pathway from ipRGCs to the rod lineage (D’Souza et al., 2024) further raises the possibility that such signalling, in addition to the canonical HIF-1α–CD73 axis, could modulate CD73 expression in developing rods. Finally, although we detected no defects in cell fate specification, both CD73-mediated and bidirectional ENT-mediated pathways can also act as sources of extracellular adenosine in the OIR model (Zhang et al., 2022).

Moreover, extracellular adenosine in the adult retina rises in darkness via a CD73-mediated pathway, whereas the circadian rhythm influences extracellular adenosine levels by changing the uptake to intracellular adenosine through ENTs (Cao et al., 2020; Ribelayga & Mangel, 2005). These observations suggest that bidirectional ENTs flux might compensate for the loss of CD73 during retinogenesis. Future studies that manipulate light exposure or circadian timing should uncover roles for CD73 in retinal development.

### Spatial CD73 expression uncovers the early astrocyte-lineage window

Pax2^+^ astrocyte progenitor cells (APCs) emerge in the optic nerve head and generate the entire retinal astrocyte population. After E16.5, some APCs differentiate centrally, whereas the rest migrate centrifugally and gradually transition into immature astrocytes (IACs) that begin to up-regulate GFAP. These early astrocyte lineages (APCs and IACs) reach the retinal margin by P3–P4, and many cells remain mitotically active until P5–P6. Consequently, the peripheral retina is preferentially derived from Pax2^+^ APCs (Chan-Ling et al., 2009; Duan et al., 2023; Duan et al., 2017; Tao & Zhang, 2014; West et al., 2005). Based on this developmental process, our spatiotemporal lineage-tracing data indicate that CD73 is transiently up-regulated in the early astrocyte lineage during migration and down-regulated once maturation is underway.

Although genetic studies using GFAP-Cre show that deleting HIF-2α does not disturb physiological vascular development (Weidemann et al., 2010), Pax2^+^/GFAP^-^astrocyte-progenitor cells (APCs) rely more on HIF-2α than on HIF-1α (Duan et al., 2014). HIF-2α becomes the dominant isoform under moderate or chronic hypoxia and drives genes encoding angiogenic and growth factors, whereas HIF-1α is preferentially stabilised during acute or severe hypoxia and orchestrates glycolytic reprogramming (Jiang et al., 2021; Sato & Yanagita, 2013; Semba et al., 2016; Yuan et al., 2024). As detailed above, CD73 is a direct HIF-1α target (Synnestvedt et al., 2002; Yuan et al., 2024), and our lineage-tracing result shows that descendants of CD73^+^ astrocytes accumulate in the peripheral retina. Collectively, our findings and earlier studies suggest that HIF-1α is active primarily within the earliest Pax2^+^/GFAP^-^ astrocyte-progenitor window and drives CD73 expression.

In addition, we find that endothelial cells lack CD73 under normoxic conditions, a result that is consistent with previous reports demonstrating that both HIF-1α and HIF-2α are inactive in endothelial cells at physiological oxygen tension but are readily induced by acute (OIR) or chronic (PHD2 deficiency) hypoxia (Duan et al., 2024; Liu et al., 2017). Future work that modulates hypoxic intensity and shifts the HIF-1α/HIF-2α balance should clarify the mechanisms governing CD73 expression during astrocyte and vascular development. Beyond oxygen signaling, APC migration and proliferation rely on PDGF-A from RGCs, whereas the APC-to-IAC transition is promoted by CNTF and additional cues from RGCs, Müller glia, and amacrine cells (Paisley & Kay, 2021; Tao & Zhang, 2014). CD73 is also known to be regulated from the transcriptional level to post-transcriptional and even post-translational levels (Alcedo et al., 2021). Therefore, these HIF-independent cues may fine-tune CD73 expression within the astrocyte lineage.

In summary, our spatiotemporal analysis across retinal progenitor and astrocyte differentiations indicates that gradients of hypoxia, retinal-wave activity, light, and adenosine metabolism may jointly regulate CD73 expression.

### Loss of CD73 disrupts normal recovery of rod photoresponses

In retinal neurons, A2A receptors are predominantly expressed in the inner retina (Chen et al., 2024; Santiago et al., 2020; Zhong et al., 2021). Although in-situ hybridization likewise shows that A2A mRNA is mainly expressed in the inner retina, only a weak signal was detected in the ONL (Kvanta et al., 1997), whereas it rises in cones during the subjective night (Li et al., 2013).

Electrophysiologically, extracellular adenosine acts in two opposite ways via A2A receptors. In dark-adapted rods, extracellular adenosine binds to A2A-like receptors on the synaptic terminal, lowers L-type Ca^2+^ influx, and thereby reduces Ca^2+^-dependent glutamate release; pharmacological blockade of these receptors does not alter the rod’s own photovoltage (Stella et al., 2003). Conversely, when adenosine acts on A2A receptors expressed by cones, extracellular adenosine increases rod–cone gap-junction coupling, thereby strengthening rod-driven cone responses and enhancing night-time cone sensitivity (Cao et al., 2020; Li et al., 2013).

By contrast, our CD73 KO mice are presumably exposed to chronically low extracellular adenosine during the subjective day. In photopic ERG, the CD73 KO showed no deficit in cone-driven responses. In contrast to the earlier pharmacological receptor blockade data, our single-cell recordings for rods revealed that genetic deletion of CD73 delayed response recovery in phototransduction.

Because ENTs mediate bidirectional adenosine flux and favour extracellular-to-intracellular uptake during the subjective day (Cao et al., 2020; Ribelayga & Mangel, 2005), we hypothesised that CD73 deficiency might reduce extracellular adenosine pools and compromise adenosine salvage, thereby limiting ATP availability in rods. Real-time ATP imaging in rods should clarify these alternative mechanisms (He et al., 2021).

### Light exposure highlights roles of CD73 in rod photoreceptors for overall retinal homeostasis

Photoreceptor apoptosis can be triggered by acute light damage or by chronic phototransduction stress (Hao et al., 2002; Wenzel et al., 2005). Unexpectedly, CD73 KO mice exhibited a thicker ONL in the inferior retina than WT, even though phototransduction stress typically leads to ONL thinning. Since the rods in the single-cell recordings were randomly selected and not regionally restricted, the measurements do not directly reflect the changes observed in this specific inferior region. We therefore propose that loss of CD73 perturbs overall retinal homeostasis, triggering changes in the ONL microenvironment that become most evident in the inferior retina. Consistent with this hypothesis, light exposure preferentially up-regulates Müller-cell-derived growth factors such as bFGF in the inferior retina (Stone et al., 1999), and CD73 deficiency impairs Müller-cell volume regulation under hypo-osmotic stress (Wurm et al., 2010). Together, our findings and previous reports suggest that CD73 activity in rods cooperates with Müller glia to preserve retinal integrity under normal lighting conditions.

### CD73 expression in the RPE is regulated independently of that in the neural retina

During development, the RPE also releases ATP into the extracellular space to modulate retinal progenitor proliferation (Pearson et al., 2005), suggesting that CD73, whose expression we identified in this study, may help maintain the extracellular ATP/adenosine balance. In adulthood, the RPE both supports the high metabolic demands of photoreceptor phototransduction and phagocytoses rod outer segments in a circadian manner (Bhoi et al., 2023; Organisciak & Vaughan, 2010). Moreover, because HIF-1α becomes active in the early phase of oxygen tension changes within the RPE (Kurihara et al., 2016), even subtle shifts in O_2_ that occur as the RPE maintains photoreceptor homeostasis during daily life may influence CD73 dynamics, thereby modulating the intra- and extracellular ATP/adenosine metabolism. In future studies, a comprehensive whole-mount analysis of the RPE will be valuable for determining the factors that shape it.

In conclusion, our study reveals that CD73 expression undergoes a spatiotemporal shift during development, and that CD73 in rods contributes the proper termination of phototransduction during adulthood. Dissecting how everyday physiological cues (e.g., hypoxia, light exposure, energy metabolism, and circadian rhythms) modulate the intrinsic dynamics of CD73 will help disentangle multiple redundant and compensatory pathways of adenosine signalling and could uncover new therapeutic targets for retinal disorders caused by lifestyle factors, such as diabetic retinopathy or age-related macular degeneration.

## Materials and methods

### Key resources table

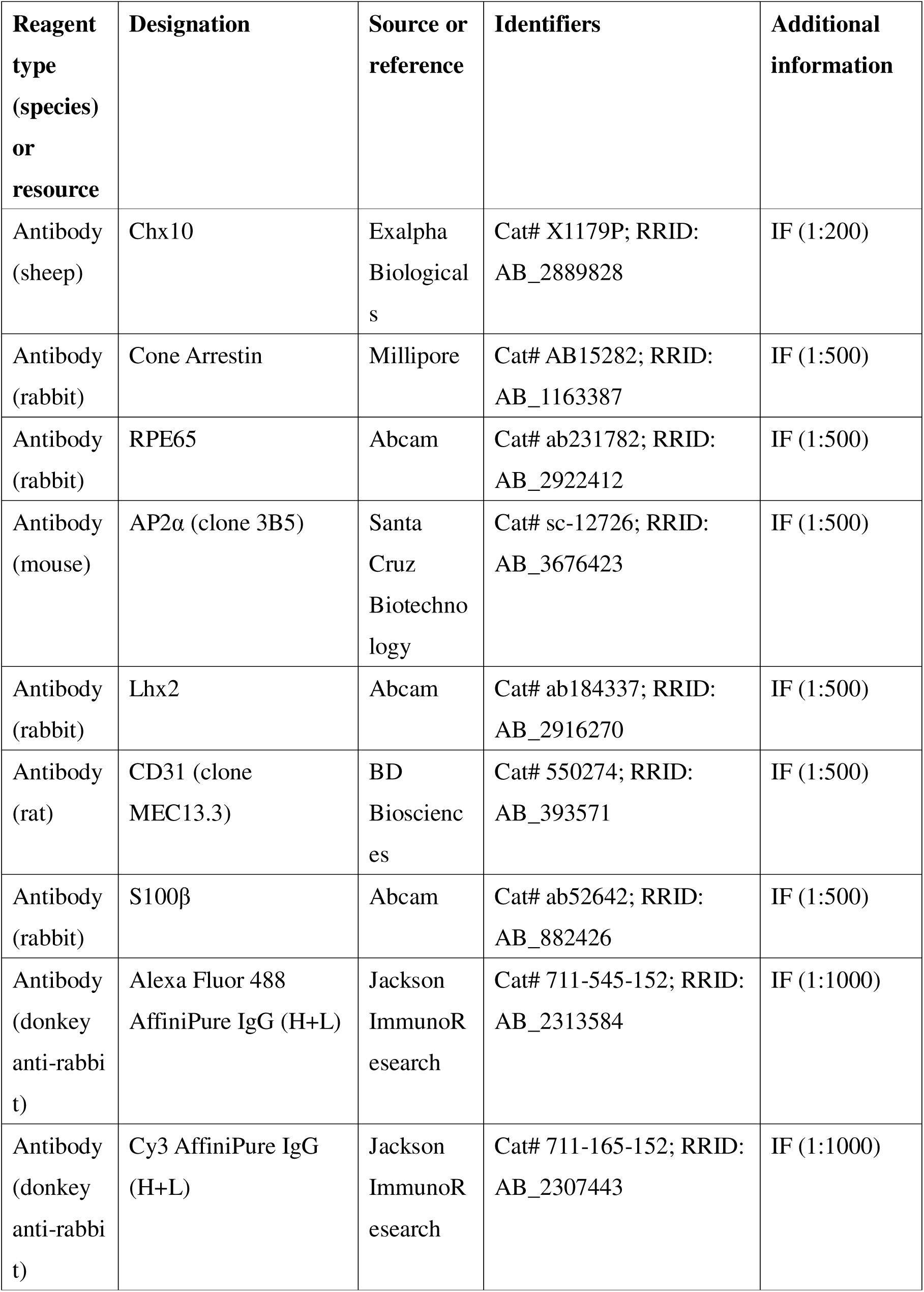
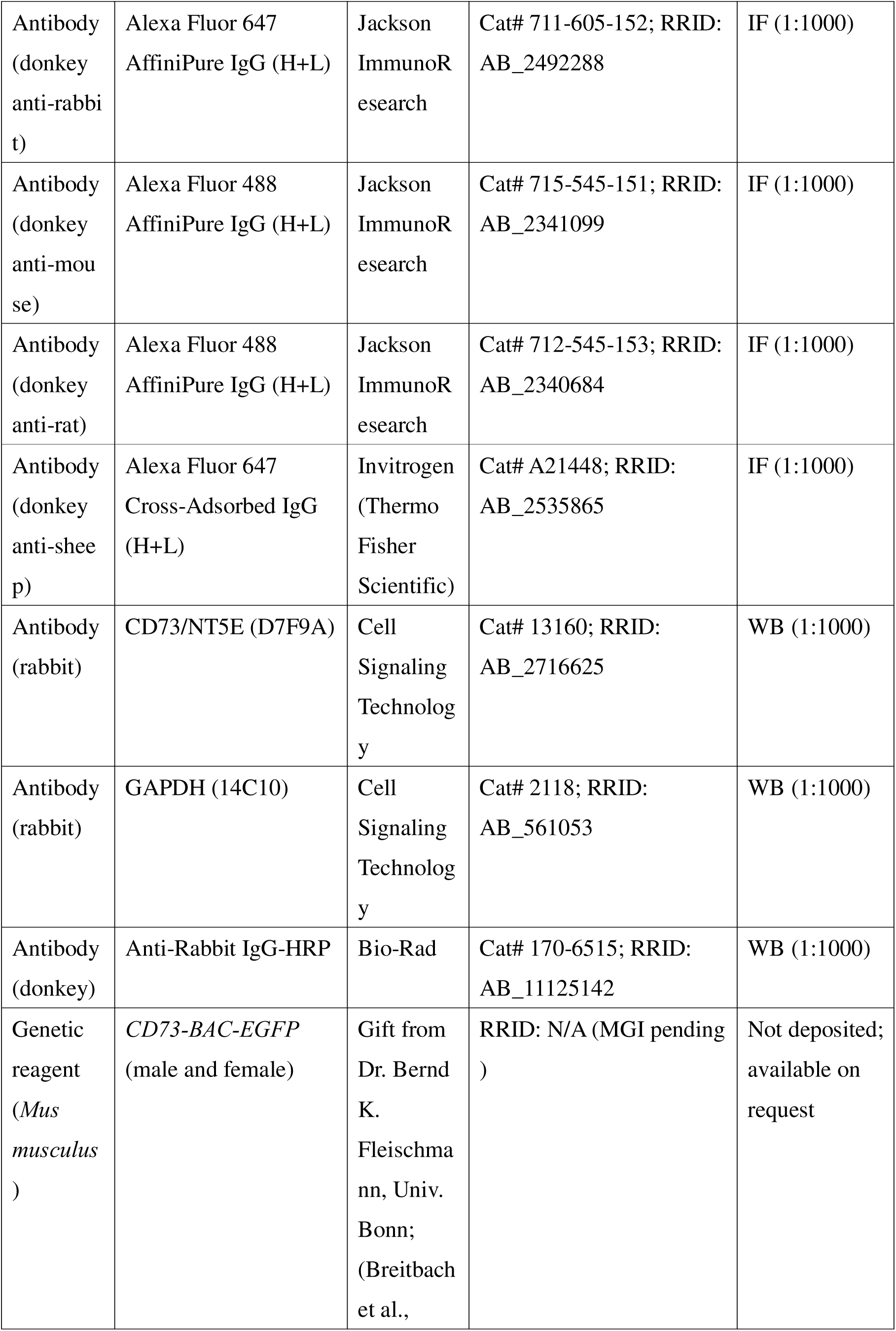
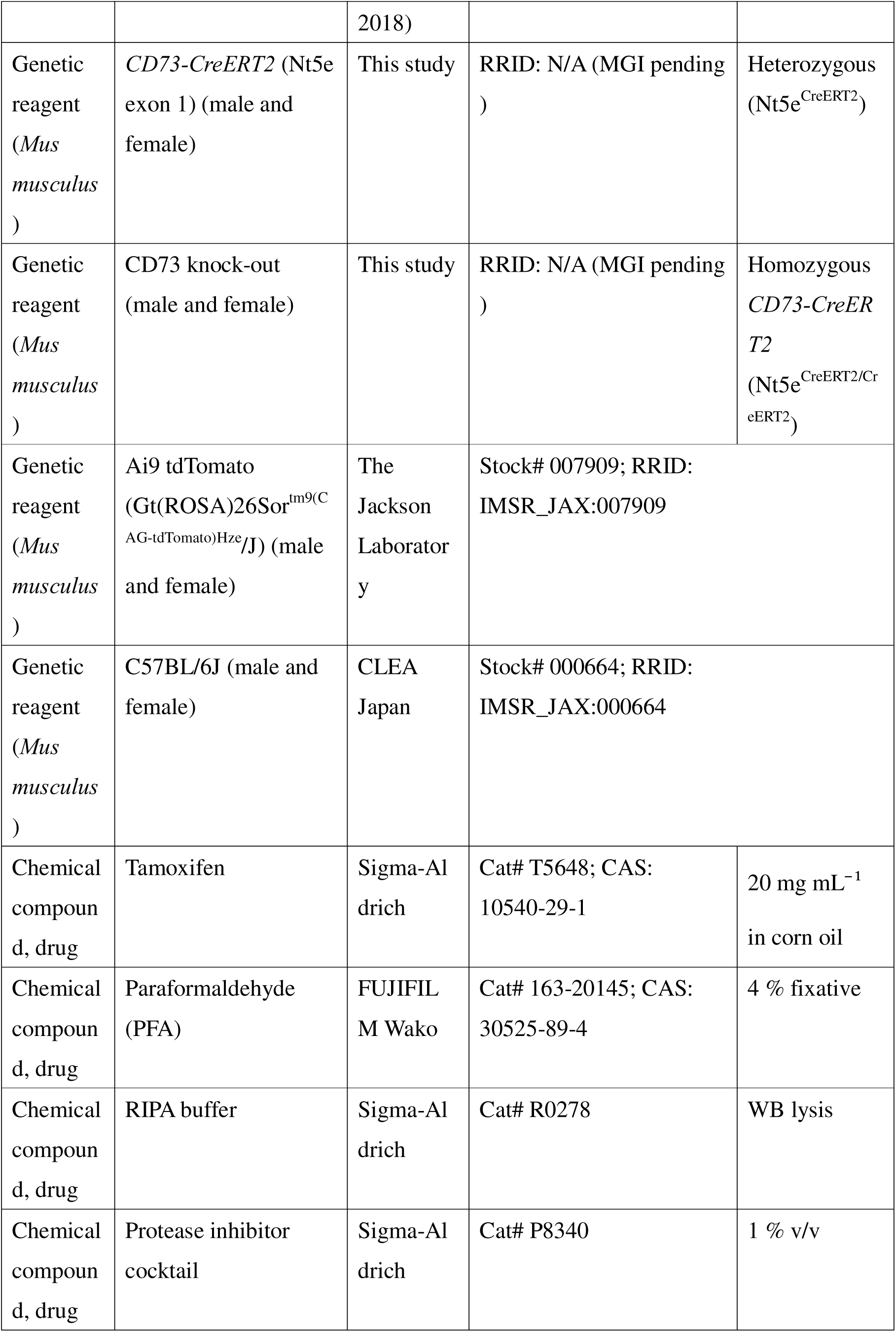
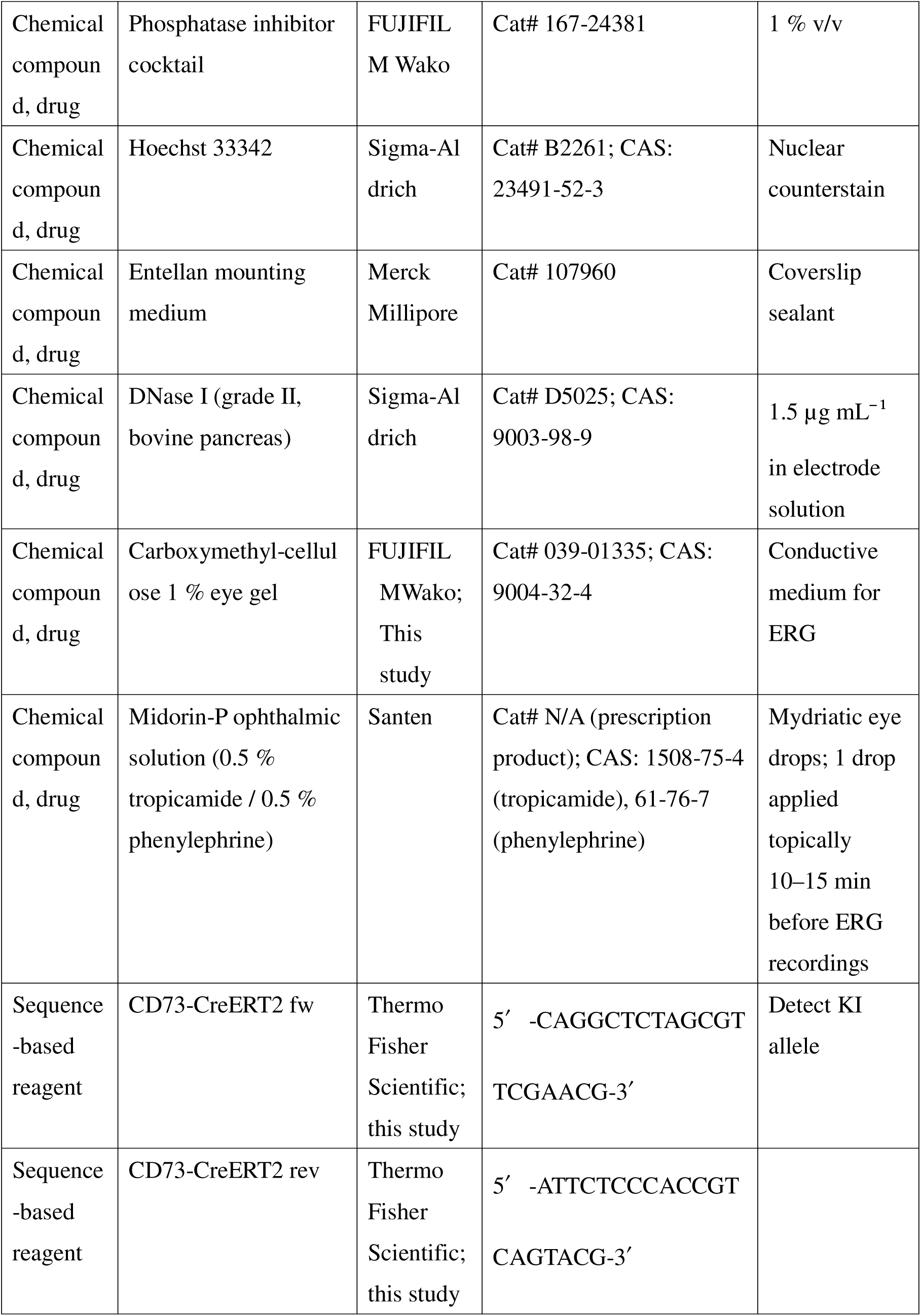
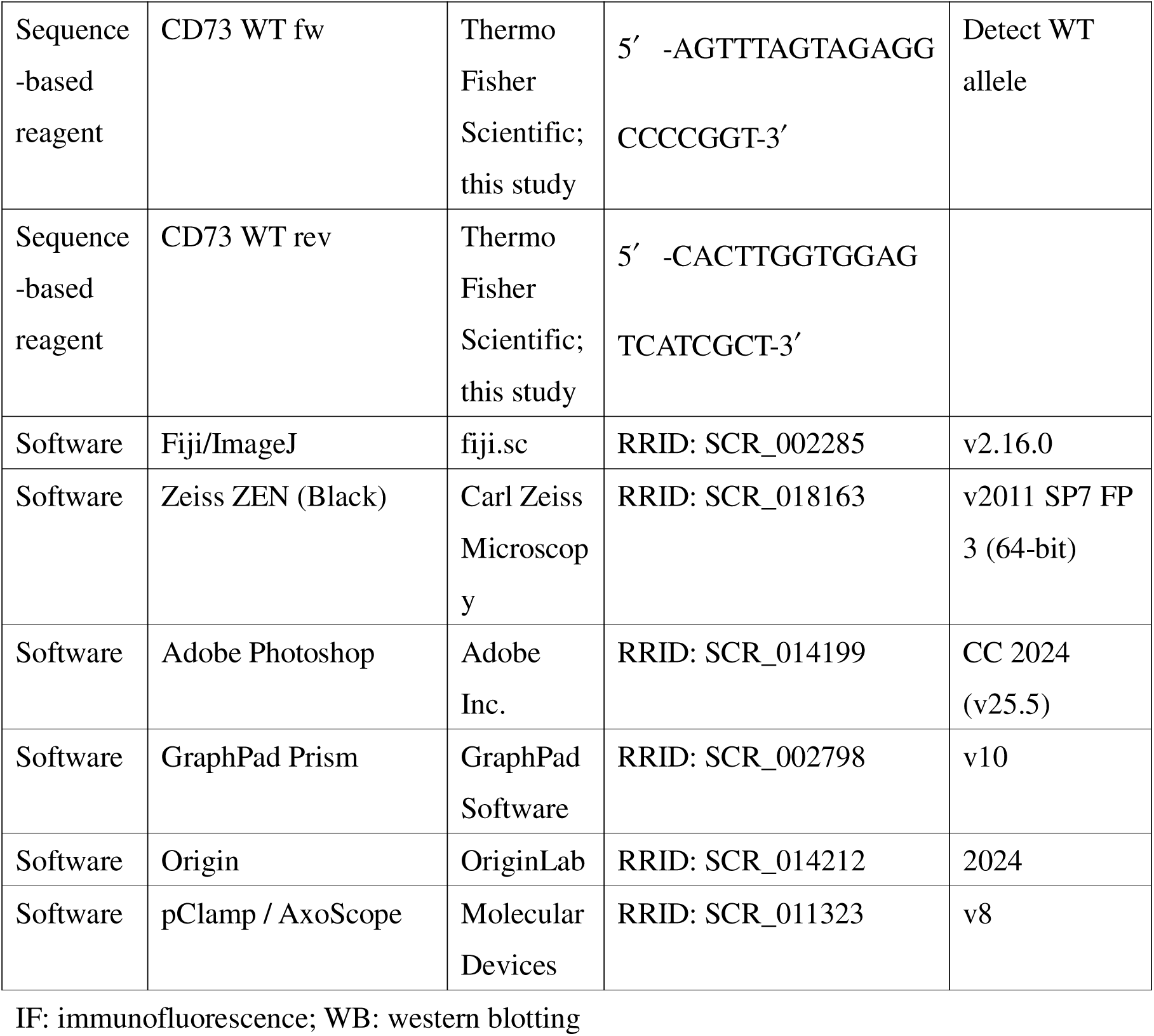

### Mice

The *CD73-BAC-EGFP* (*CD73-EGFP*; RRID: N/A) mouse line was previously described and generously provided by Dr. Bernd K. Fleischmann (University of Bonn, (Breitbach et al., 2018)). To generate the *CD73-CreER^T2^* knock-in mouse line (*Nt5e*^CreERT2^; RRID: N/A), a CreERT2 cassette was inserted into exon 1 of the mouse Nt5e locus using a CRISPR/Cas9-based genome editing platform. CD73 KO mice were generated by crossing *CD73*-*CreER^T2^* knock-in mice as homozygotes. For lineage tracing experiments, *CD73-CreER^T2^* mice were crossed with Rosa26-loxp-STOP-loxp-tdTomato (tdTomato) knockin mice (C57BL/6J; Jackson Laboratory, #007909; RRID: IMSR_JAX:007909). In this lineage-tracing experiment, the CreERT2 cassette is inserted into one allele of Nt5e, resulting in heterozygous expression of CD73. Wild-type C57BL/6J mice (Stock # 000664; RRID: IMSR_JAX:000664) were purchased from CLEA Japan, Inc. All mice were bred and maintained under a 12-hour light/dark cycle (lights on at 7:00 A.M. [ZT0], lights off at 7:00 P.M. [ZT12]) and used for experiments between ZT3 (10:00 A.M.) and ZT9 (4:00 P.M.). Both male and female mice were used, and data from both sexes were pooled because no sex-specific differences were detected. All procedures were approved by the Animal Care and Use Committee of the University of Tsukuba and complied with institutional and governmental guidelines (approved number: 23-432). Mice were euthanized by intraperitoneal injection of a triple anesthetic mixture (8 mg/kg midazolam, 1.5 mg/kg medetomidine, and 10 mg/kg butorphanol tartrate, or higher if required), followed by cervical dislocation to ensure complete euthanasia.

### Mouse genotyping

Genomic DNA was extracted from mouse tail biopsies and used as a template for PCR-based genotyping. Two sets of primers were employed to detect the presence of either the *CD73-CreER^T2^* knock-in allele or the WT CD73 allele. Primer pairs (Thermo Fisher Scientific; this study) were: *CD73-CreER^T2^*knock-in (fw: 5′-CAGGCTCTAGCGTTCGAACG-3′, rev: 5′-ATTCTCCCACCGTCAGTACG-3′; product size: 182 bp) and CD73 WT (fw: 5′-AGTTTAGTAGAGGCCCCGGT-3′, rev: 5′-CACTTGGTGGAGTCATCGCT-3′; product size: 235 bp).

### Tamoxifen treatment

For lineage tracing, 1- to 2-month-old *CD73-CreER^T2^* mice (both sexes) received tamoxifen (2 mg per mouse per day; Sigma-Aldrich; 20 mg/mL in corn oil) via intraperitoneal injection for three consecutive days. Neonatal mice (≤14 days old) received a single oral dose of tamoxifen (100 µg/g body weight). For embryonic-stage analyses, pregnant dams received a single 2 mg oral dose of tamoxifen. All tamoxifen-treated mice were raised to 3 months of age before euthanasia. To assess potential Cre “leakiness,” we also analyzed *CD73-CreER^T2^;Rosa-tdTomato* mice that did not receive tamoxifen.

### Staining of retinal sections

Enucleated eyes were fixed in 4% (w/v) paraformaldehyde (PFA) for 45 min at room temperature, then rinsed in PBS. Fixed tissues were cryoprotected overnight in 30% (w/v) sucrose in PBS at 4°C, embedded in OCT compound (Tissue-Tek, Sakura), and rapidly frozen. Retinal sections (12 µm) were cut on a cryostat and air-dried before staining.

For hematoxylin and eosin (H&E) staining, sections were briefly rinsed in distilled water, stained with hematoxylin (Wako #131-09665) for 5 min, and counterstained with eosin Y (Wako #058-00062) for 45 s. After dehydration in a graded ethanol series, sections were mounted with Entellan (Merck Millipore).

For immunostaining, sections were blocked in PBS containing 3% donkey serum and 0.3% Triton X-100 for 1 h at room temperature. Primary antibodies, diluted in the same blocking solution, were applied overnight at 4°C. After washing in PBS, sections were incubated for 1 h at room temperature with Alexa Fluor–conjugated secondary antibodies. Hoechst 33342 (Sigma-Aldrich #B2261) was added at 1:1000 to counterstain nuclei. Slides were then washed and coverslipped. Antibody information is listed in Key resources table. Images were captured with a Zeiss LSM700 confocal or a Zeiss Axio Imager.Z2 widefield microscope. Brightness and contrast were adjusted uniformly in Adobe Photoshop.

### Whole-mount immunostaining

Eyes were fixed in 4% PFA for 10 min at room temperature. Retinas were then isolated and post-fixed in 4% PFA for 1.5 h at 4°C. After two PBS washes, retinas were blocked in 3% donkey serum and 0.3% Triton X-100 in PBS for 2 h at room temperature before incubation with primary antibodies for five days at 4°C. Retinas were washed in PBS and incubated overnight at 4°C with secondary antibodies and Hoechst. Finally, retinas were washed, radially cut, and mounted flat with antifade medium. Antibody information is listed in Key resources table. Images were captured with a Zeiss LSM700 confocal or a Zeiss Axio Imager.Z2 widefield microscope. Confocal images are shown as single optical sections or z-stack projections.

All images shown are representative of at least three biological replicates (n = 3 mice per time point).

### Western blot analysis

Retinas were harvested from 2–3-month-old mice, snap-frozen in liquid nitrogen, and minced with a needle. Tissues were then lysed in RIPA buffer (Sigma-Aldrich, #R0278) supplemented with 1% phosphatase inhibitor cocktail (Wako, #167-24381) and 1% protease inhibitor cocktail (Sigma-Aldrich, #P8340). The lysates were mixed with 3× SDS sample buffer containing 2-mercaptoethanol, boiled at 95°C for 5 min, and subjected to SDS-PAGE. Proteins were then transferred to a polyvinylidene difluoride (PVDF) membrane (Millipore, #IPHV00010), blocked, and immunoblotted with the following primary antibodies: NT5E/CD73 (D7F9A) rabbit mAb (1:1000, Cell Signaling, #13160) and GAPDH (14C10) rabbit mAb (1:1000, Cell Signaling, #2118). Membranes were then incubated with anti-rabbit HRP-conjugated secondary antibodies (1:1000, Bio-Rad, #170-6515; RRID: AB_11125142) for 1 h at room temperature. For visualization, a chemiluminescence kit (Atto, #2332637) was used, and signals were detected using a WSE-6300 LuminoGraph III (Atto).

### Histological quantification

#### Quantification of CD73CreER^T2^-labeled cell density

For lineage tracing experiments, mice (n ≥ 3 per group) received tamoxifen at various time points ranging from E13.5 to 2 months of age (see “Results” for details) and were all sacrificed at 3 months of age. Whole-mount retinas were imaged using a confocal microscope (10× objective, 0.5× digital zoom). To avoid overlap between central and peripheral regions, four central fields (near the optic nerve) and five peripheral fields (near the retinal edge) were captured per retina. In order to exclude the ONL, maximum-intensity z-projections were generated only from the INL to the GCL.

For quantification analysis, each of the four central fields was divided into two zones: 0–500 µm (C1) and 500–1000 µm (C2) from the optic nerve head. Likewise, each of the five peripheral fields was divided into two zones: 0–500 µm (P2) and 500–1000 µm (P1) from the retinal edge. In each zone of each field, tdTomato^+^ astrocytes and tdTomato^+^ cells in the INL were manually counted using the “Cell Counter” tool in FIJI v2.16.0 (RRID: SCR_002285; see Key resources table). The area of each zone was measured in the same software using the freehand selection tool. The density of tdTomato^+^ cells was then calculated as the number of cells per 0.1 mm². Because multiple fields were obtained per mouse, the densities and area measurements from the same zone type (e.g., all C1 fields in one retina) were averaged to yield a single zone-based value per zone per mouse (i.e., one value for C1 per mouse, one for C2, etc.). For temporal analysis, these zone-averaged densities were averaged across mice at each tamoxifen administration time point, and, for spatial analysis, we plotted the zone-based densities (C1, C2, P1, P2) for each tamoxifen injection time point.

All statistical analyses were performed in GraphPad Prism v10 (RRID: SCR_002798; see Key resources table). One-way ANOVA was used to compare tdTomato^+^ cell densities among different time points (for temporal analysis) and among zones (C1, C2, P1, P2; for spatial analysis), followed by Tukey’s multiple comparisons test. Significance was defined as *p* < 0.05. The exact sample size (n) is indicated in each figure legend, and results are expressed as mean ± s.e.m., with each dot in the graphs representing an individual mouse.

#### Cell genesis quantification

Retinal cell genesis was examined at P14 using a modified protocol (Zhang et al., 2023). Briefly, 20-µm sections were prepared and immunolabeled with specific markers for retinal cell types (see “Results” for details). Three sections per mouse were imaged (20× objective, 0.8× digital zoom; field size, 400.1 µm x 400.1 µm) in both the superior and inferior retina at 500–1000 µm from the optic nerve. Cell counts were averaged for each mouse. All statistical analysises were performed using GraphPad Prism v10 (GraphPad Software), and exact n numbers are provided in the figure legends. Normality was assessed with the Shapiro-Wilk test, and groups were compared using unpaired two-tailed Student’s *t*-test (*p* < 0.05 was considered statistically significant).

#### Retinal morphometry

Retinal sections were obtained from WT and CD73 KO mice at P14, 1 month, and 3 months of age (n ≥ 7 per group). Sections were cut along the vertical meridian through the optic nerve and stained with H&E. ONL thickness was measured in 250-µm increments from the optic nerve head toward the retinal periphery (superior and inferior) using FIJI. Retinal morphometry graphs were generated using Origin 2024 (OriginLab). Statistical analyses were performed with GraphPad Prism v10 (GraphPad Software), and exact n numbers are provided in the figure legends. Normality was assessed with the Shapiro-Wilk test, and groups were compared using Student’s *t*-test (*p*-value < 0.05 was considered statistically significant). Data are shown as mean ± s.e.m.

### in vivo ERG

Mice at 2-3 months of age were dark-adapted overnight and the following procedures were performed under dim red light. The animals were anesthetized with midazolam (4 mg/kg), medetomidine (0.75 mg/kg), and butorphanol tartrate (5 mg/kg) before the recordings as previously described (Miwa et al., 2019). The animal’s pupils were dilated with eye drops of 0.5% tropicamide and 0.5% phenylephrine. The anesthetized animal was mounted on a stage with a heat pad inside a light-proof faraday cage. A looped platinum recording electrode was placed on the cornea with a drop of 1.0% carboxymethyl cellulose and a reference electrode was positined in the oral cavity. A ground electrode was clipped to the tail. Full-field light stimulus was delivered from a custom-made Ganzfeld dome. The stimulus, emitted from a green LED (M505L4, Thorlab) was attenuated using neutral density filters and transmitted to the Ganzfeld dome via a liquid light guide. For the photopic ERG, the mouse eye was light adapted with background white light (30 cd/cm^2^). All mice were kept warm during the experiment using a heat pad. The photovoltage was amplified and filtered using a differential amplifier (bandpass: 0.5 to 300 Hz, Nihon Koden) and digitized at 10 k Hz with Digidata1320A and AxoScope8. To analyze the amplitude of a- and b-wave, averaged responses were lowpass filtered below 70 Hz to eliminate oscillatory response beforehand. The implicit time (time to reach the peak amplitude) of the b-wave response evoked by dim light was estimated by fitting the response with the Gaussian function; *r(t) =R*· 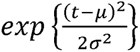, where *R* is the peak amplitude of the response, μ is the implicit time, and σ is the Gaussian width. The amplification constant of rod photoreceptorswas estimated by fitting the initial phase of the a-wave with the equation proposed by Pugh and Lamb (Pugh & Lamb, 1993)

### Rod Single-Cell Recordings

For the single-cell recordings, the light-dependent current of rod photoreceptors was recorded by drowning the outer segments into a suction pipette as previously described (Sakurai et al., 2007). Briefly, a mouse aged 2-3 months was dark-adapted before the experiment. All the following procedures were performed under infrared light. The animals were euthanized by cervical dislocation and the dissociated retina was stored in equilibrated Locke’s solution (112 mM NaCl, 3.6 mM KCl, 2.4 mM MgCl_2_, 1.2 mM CaCl_2_, 10 mM HEPES, 20 mM NaHCO_3_, 3mM Na_2_-succinate, 0.5 mM Na-glutamate 10 mM glucose, 0.1% MEM vitamin, and 0.1% MEM non-essential amino acids) before use at RT. The recording chamber was perfused with Locke’s solution equilibrated with 95%O_2_/5%CO_2_ gas and preheated to 32-36℃. The recording electrode with an inter diameter adjusted to the width of the outer segment was prepared and filled with an electrode solution (140 mM NaCl, 3.6 mM KCl, 2.4 mM MgCl_2_, 1.2 mM CaCl_2_, 3 mM HEPES, and 0.02 mM EDTA at pH 7.4). A piece of retina chopped with a razor blade in the electrode solution with 1.5 μg/mL DNaseI was transferred to the recording chamber mounted on the inverted microscope stage (IX73, Olympus). The photocurrent signal was amplified with Axopatch 200B amplifier, low-pass filtered with a cutoff frequency of 30 Hz, and digitized at 1 k Hz using Digidata1322A and pClamp8 (RRID: SCR_011323; see Key resources table). The sensitivity (*I*_1/2_) of photoreceptor was estimated by fitting the intensity-response data for each photoreceptor to the equation: 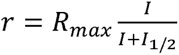, where *R_max_* is the maximal response amplitude (pA), *I* is the flash intensity, and *I*_1/2_ is the sensitivity (photons/μm^2^).

### Statistical analysis

Statistical analyses were performed using GraphPad Prism v10 (GraphPad Software). All data are reported as the mean ± s.e.m. Normality was assessed using the Shapiro-Wilk test. For comparisons between two groups, an unpaired two-tailed Student’s *t*-test was used. For comparisons among three or more groups, one-way ANOVA was performed, followed by Tukey’s multiple comparisons test. Significance was defined as *p* < 0.05.

## Data availability

All data generated or analysed during this study are included in the manuscript and supporting files.

## Acknowledgements

We thank Dr. Satoru Takahashi and Dr. Erna Raja for their experimental support, Dr. Fuminori Tsuruta, Dr. Yuya Sanaki, and Dr. Masafumi Muratani for critically reading the manuscript. We are also grateful to the Laboratory Animal Resource Center at the University of Tsukuba for their professional care of the animals, as well as to Ms. Mariko Higashi, Ms. Mami Ho, Ms. Tomomi Zama, and Ms. Eri Motoyama for their technical assistance. This study was supported in part by JSPS KAKENHI Grant Numbers JP22K16940 (to R.I.) and JP22K09346 (to K.K.), the TERUMO Life Science Foundation, the Uehara Memorial Foundation, the NOVARTIS Foundation, the Inamori Foundation, SENSHIN Medical Research Foundation, Japan Agency for Medical Research and Development Grant Numbers 23bm1123032h0001.

## Author Contributions

Conceptualization: R.I.; Formal analysis: R.I., K.S.; Investigation: R.I., K.S., N.H., S.M., K.K.; Writing – original draft: R.I., K.S.; Writing – review & editing: R.I., K.S., B.K.F., S.M., K.K., H.Y.; Supervision: R.I., H.Y.; Project administration: R.I.; Funding acquisition: R.I., K.K.

## Figure legends

***Figure 1—figure supplement 1.*** Retinal layer formation from P1 to P14.

This schematic shows the developmental progression of mouse retinal layers from birth (P1) through P14. At P1, the retina comprises two nuclear layers: the GCL and the NBL. By approximately P3, the NBL divides into an outer (oNBL) and inner (iNBL) neuroblastic layer, and the outer plexiform layer (OPL) first becomes detectable. From around P5–P6 onward, the oNBL and iNBL are referred to as the outer nuclear layer (ONL) and the inner nuclear layer (INL).

Abbreviations: GCL, ganglion cell layer; iNBL, inner neuroblastic layer; INL, inner nuclear layer; NBL, neuroblastic layer; oNBL, outer neuroblastic layer; ONL, outer nuclear layer; OPL, outer plexiform layer.

***Figure 1—figure supplement 2***. CD73-EGFP expression pattern at P0 and P1 in the whole-mount retinas.

Representative images of wholemount from CD73-EGFP mice at postnatal day 0 (P0, left) and P1 (right). Orthogonal views are shown for the xy plane (center), xz plane (top), and yz plane (right). The white lines in the xz and yz views mark the z-position of the xy plane. The green and magenta lines indicate the y- and x-positions, respectively, which were used to generate the cross-sectional planes.

Abbreviations: GCL, ganglion cell layer; NBL, neuroblastic layer.

Scale bars: 20 µm.

***Figure 1—figure supplement 3.*** CD73-EGFP expression is detected in the rod lineage from P5 to P14.

(A) Representative images of transverse sections from P5 to P14. Arr3 (cone arrestin, magenta) marks the cone lineage.

(B) Higher-magnification view of the dashed box in (A). The top panels show the Arr3 (magenta) channel, while the bottom panels show the CD73-EGFP (green) channel.

Scale bars: 50 µm (A), 25 µm (B).

***Figure 2—figure supplement 1.*** *CD73-CreER^T2^;tdTomato* lineage-tracing reveals CD73 expression in the RPE as well as in the ONL of adult retinas.

(A) Representative images from *CD73-CreER^T2^* mice without tamoxifen injection (no-injection control). The left panel shows a whole-mount view of the retina, and the right panel is a cross-sectional view. Left panel: CD31 (green) labels blood vessels. Right panel: Hoechst (green) labels nuclei. tdTomato is shown in magenta in both panels.

(B) Representative femoral bone image from *CD73-EGFP; CD73CreER^T2^;tdTomato* mice treated with tamoxifen at 3 months of age. EGFP+ (green)/tdTomato+(red) cell was observed at the inner surface of cortical bone. Hoechst (blue) labels nuclei.

(C) Immunostaining of retinal sections from *CD73-CreER^T2^* mice two weeks after tamoxifen injection. The left panel shows a low-magnification view of the entire retina, with RPE65 (green) labeling the RPE. tdTomato^+^ cells (magenta) are visible in the RPE layer. The right panels show higher-magnification views of the white dashed area in left panel. The top row is the merged image, the middle row is RPE65 alone, and the bottom row is tdTomato alone.

Abbreviations: BM, bone marrow; CB, cortical bone; ONL, outer nuclear layer; RPE, retinal pigment epithelium.

Scale bars: 500 µm (A, left); 100 µm (A, right; C, left); 25 µm (C, right); 10 µm (B).

***Figure 2—figure supplement 2.*** *CD73-CreER^T2^* lineage tracing demonstrates CD73^+^ cells in the RPE, INL, and RNFL as well as the rod lineage within ONL during retinal development.

(A) Distinct sets of *CD73-CreER^T2^* mice were each treated with tamoxifen at P10, P14, 1 month, or 2 months of age, and all were examined at 3 months of age. Nuclei are labeled with Hoechst (green), and tdTomato is shown in magenta. Arrowheads indicate tdTomato^+^ cells in the RPE, arrows mark tdTomato^+^ cells in the INL.

(B) Higher-magnification views of the outer portion of the ONL (cone-rich region), in the same groups described in (A). Arr3 (cone arrestin, green) labels cone photoreceptors; tdTomato is shown in magenta. The top rows show merged images, the middle rows show Arr3 alone, and the bottom rows show tdTomato alone.

(C) Distinct sets of *CD73-CreER^T2^* mice were each treated with tamoxifen at E13.5, E14.5, E15.5, E16.5, and E17.5, and all were examined at 3 months of age. Nuclei are labeled with Hoechst (green), and tdTomato (magenta) is visible in the RPE (arrowheads).

(D) Immunostaining of retinal sections from a *CD73-CreER^T2^* mouse treated with tamoxifen at P3 and analyzed at 3 months of age. Nuclei are labeled with Hoechst (green), and tdTomato is shown in magenta. Arrowheads indicate tdTomato^+^ cells, presumed to reside in the RNFL immediately vitreal to the GCL.

Abbreviations: GCL, ganglion cell layer; INL, inner nuclear layer; ONL, outer nuclear layer; RNFL, retinal nerve fiber layer; RPE, retinal pigment epithelium.

Scale bars: 100 µm (A; C; D); and 25 µm (B).

***Figure 3—figure supplement 1.*** Representative images illustrating the anatomy of the adult eye.

(A) An H&E-stained cross-section of the adult eye, showing the spatial characteristics of each compartment.

(B) A whole-mount immunostaining of the adult retina with CD31 (a vascular endothelial marker) shown in green.

Scale bars: 500 µm (A); 1000 µm (B).

***Figure 4—figure supplement 1.*** Descendants of *CD73-CreER^T2^*^+^ cells in the astrocyte lineage in the central retina.

(A) Representative wholemount retinal images from *CD73-CreER^T2^;tdTomato* mice treated with tamoxifen at E16.5, E17.5, E18.5, or P3. Shown are maximum-intensity projection images spanning from the INL to the RNFL, excluding the ONL, giving an overview of the central retina in each tamoxifen treatment group. Nuclei are labeled with Hoechst (green), and tdTomato is shown in magenta.

(B) Higher-magnification views of the white dashed boxes in (A). The astrocyte marker S100β is labeled in green, and tdTomato is shown in magenta. Arrowheads indicate tdTomato and S100β double-positive cells.

Abbreviations: GCL, ganglion cell layer; INL, inner nuclear layer; ONL, outer nuclear layer; RNFL, retinal nerve fiber layer.

Scale bars: 500 µm (A); 50 µm (B).

***Figure 5—figure supplement 1.*** The distribution pattern of descendants of *CD73-CreER^T2+^* cells in the INL cells of the peripheral retina.

(A) Representative wholemount retinal images from *CD73-CreER^T2^;tdTomato* mice treated with tamoxifen at P7, P10, P14, or 1 month of age. Shown are maximum-intensity projection images from the INL to the RNFL, excluding the ONL, providing an overview of the peripheral retina for each treatment group. tdTomato is shown in magenta. Arrowheads indicate tdTomato^+^ cells in the INL.

(B) Higher-magnification views of the white dashed boxes in (A). The astrocyte marker S100β is labeled in green, and tdTomato is shown in magenta. These images confirm that the tdTomato^+^ cells in (A) reside in the INL and lack S100β expression.

Abbreviations: INL, inner nuclear layer; ONL, outer nuclear layer; RNFL, retinal nerve fiber layer.

Scale bars: 500 µm (A); 50 µm (B).

***Figure 5—figure supplement 2.*** Confirmation of CD73 expression deficiency in CD73 KO mice.

(A) Representative genotyping results. The left panel shows the *CD73-CreER^T2^* knock-in allele (primer product, 182 bp), and the right panel shows WT CD73 allele (primer product, 235 bp). In the left panel, the CD73 KO displays a band corresponding to the *CD73-CreER^T2^* knock-in allele, whereas the WT does not. In the right panel, the CD73 KO lacks the band for the WT CD73 allele, while the WT displays that band.

(B) Representative immunoblot. The expected molecular weights of CD73 (∼70 kDa) and GAPDH (∼37 kDa) are indicated. Both CD73 KO and WT samples show a GAPDH band at ∼37 kDa, but only the WT shows a CD73 band at ∼70 kDa; the CD73 KO lacks this band, confirming the deficiency of CD73 expression.

Abbreviations: bp, base pairs; KO, knockout; WT, wild-type.

***Figure 5—figure supplement 3.*** CD73 KO mice show no abnormalities in retinal cell genesis under physiological conditions.

(A) Representative retinal sections from WT and CD73 KO mice at postnatal day 14 (P14). Sections were immunostained for AP2α (amacrine cells), Vsx2 (bipolar cells), and Lhx2 (Müller glia).

(B) Quantification of AP2α^+^, Vsx2^+^, and Lhx2^+^ cells in the INL. Data are shown as mean ± s.e.m., with each dot representing an individual sample (n = 6 WT, n = 8 CD73 KO). Statistical significance was determined using unpaired two-tailed Student’s *t*-test.

(C) Measurement of ONL thickness (in µm) along the superior (0–2.25 mm) and inferior (–2.25–0 mm) hemiretina in WT and KO mice. Data are shown as mean ± s.e.m. (n = 9 WT, n = 9 CD73 KO). All statistical analyses were performed in GraphPad Prism v10 (GraphPad Software).

Abbreviations: GCL, ganglion cell layer; INL, inner nuclear layer; KO, knockout; ONL, outer nuclear layer; WT, wild-type.

Scale bars: 100 µm (A).

***Figure 6—figure supplement 1.*** Activation processes of the rod phototransduction cascade

(A) The average a-wave of WT (green, n = 10) and CD73 KO (magenta, n = 10) mice.

(B) Plot of the amplification constant of rod photoreceptors as a function of the number of the rhodopsin photoisomerizations. Data are shown as mean ± s.e.m. (n = 10 WT, n = 10 CD73 KO).

Statistical significance was assessed using unpaired two-tailed Student’s *t*-test (\**p* < 0.05).

Abbreviations: KO, knockout; WT, wild-type.

***Figure 7—figure supplement 1.*** ONL thickening in the most light-exposed region becomes apparent during adulthood.

(A) ONL thickness (µm) measured along the superior (0–2.25 mm) and inferior (−2.25–0 mm) hemiretina in WT (green) and CD73 KO (magenta) mice at 1-month-old (dashed lines) and 3-month-old (solid lines). Data are presented as mean ± s.e.m. (1-month-old: n = 11 WT, n = 7; CD73 KO; 3-month-old: n = 10 WT; n = 9 KO). Statistical significance was assessed using unpaired two-tailed Student’s *t*-test (\**p* < 0.05, \*\**p* < 0.01).

(B) Representative hematoxylin and eosin (H&E)–stained sections of WT and CD73 KO retinas at 3 months of age. Retinal sections were cut along the vertical meridian at the level of the optic nerve; the image shown is located approximately 1000 µm inferior to the optic nerve.

Abbreviations: KO, knockout; ONL, outer nuclear layer; WT, wild-type.

Scale bars: 100 µm (B).

## References

Alcedo, K. P., Bowser, J. L., & Snider, N. T. (2021). The elegant complexity of mammalian ecto-5’-nucleotidase (CD73) [Review]. Trends Cell Biol, 31(10), 829–842. 10.1016/j.tcb.2021.05.008

Alves, C. H., Pellissier, L. P., & Wijnholds, J. (2014). The CRB1 and adherens junction complex proteins in retinal development and maintenance. Progress in Retinal and Eye Research, 40, 35–52. 10.1016/j.preteyeres.2014.01.001

Bhoi, J. D., Goel, M., Ribelayga, C. P., & Mangel, S. C. (2023). Circadian clock organization in the retina: From clock components to rod and cone pathways and visual function [Review]. Prog Retin Eye Res, 94, 101119. 10.1016/j.preteyeres.2022.101119

Biswas, S., Cottarelli, A., & Agalliu, D. (2020). Neuronal and glial regulation of CNS angiogenesis and barriergenesis [Review]. Development, 147(9). 10.1242/dev.182279

Biswas, S., Shahriar, S., Bachay, G., Arvanitis, P., Jamoul, D., Brunken, W. J., & Agalliu, D. (2024). Glutamatergic neuronal activity regulates angiogenesis and blood-retinal barrier maturation via Norrin/beta-catenin signaling. Neuron, 112(12), 1978–1996 e1976. 10.1016/j.neuron.2024.03.011

Bonezzi, P. J., Stabio, M. E., & Renna, J. M. (2018). The Development of Mid-Wavelength Photoresponsivity in the Mouse Retina. Curr Eye Res, 43(5), 666–673. 10.1080/02713683.2018.1433859

Breitbach, M., Kimura, K., Luis, T. C., Fuegemann, C. J., Woll, P. S., Hesse, M., Facchini, R., Rieck, S., Jobin, K., Reinhardt, J., Ohneda, O., Wenzel, D., Geisen, C., Kurts, C., Kastenmüller, W., Hölzel, M., Jacobsen, S. E. W., & Fleischmann, B. K. (2018). In Vivo Labeling by CD73 Marks Multipotent Stromal Cells and Highlights Endothelial Heterogeneity in the Bone Marrow Niche. Cell Stem Cell, 22(2), 262–276.e267. 10.1016/j.stem.2018.01.008

Brzezinski, J. A., & Reh, T. A. (2015). Photoreceptor cell fate specification in vertebrates [Review]. Development, 142(19), 3263–3273. 10.1242/dev.127043

Burger, C. A., Jiang, D., Mackin, R. D., & Samuel, M. A. (2021). Development and maintenance of vision’s first synapse [Review]. Dev Biol, 476, 218–239. 10.1016/j.ydbio.2021.04.001

Cameron, M. A., Morley, J. W., & Pérez-Fernández, V. (2020). Seeing the light: different photoreceptor classes work together to drive adaptation in the mammalian retina. Current Opinion in Physiology, 16, 43–49. 10.1016/j.cophys.2020.05.003

Cao, J., Ribelayga, C. P., & Mangel, S. C. (2020). A Circadian Clock in the Retina Regulates Rod-Cone Gap Junction Coupling and Neuronal Light Responses via Activation of Adenosine A(2A) Receptors. Front Cell Neurosci, 14, 605067. 10.3389/fncel.2020.605067

Caprara, C., Thiersch, M., Lange, C., Joly, S., Samardzija, M., & Grimm, C. (2011). HIF1A Is Essential for the Development of the Intermediate Plexus of the Retinal Vasculature. Investigative Opthalmology & Visual Science, 52(5). 10.1167/iovs.10-6222

Chan-Ling, T., Chu, Y., Baxter, L., Weible Ii, M., & Hughes, S. (2009). In vivo characterization of astrocyte precursor cells (APCs) and astrocytes in developing rat retinae: differentiation, proliferation, and apoptosis. Glia, 57(1), 39–53. 10.1002/glia.20733

Chen, S., Zhou, S., Zang, K., Kong, F., Liang, D., & Yan, H. (2014). CD73 expression in RPE cells is associated with the suppression of conventional CD4 cell proliferation. Exp Eye Res, 127, 26–36. 10.1016/j.exer.2014.05.008

Chen, X., Sun, X., Ge, Y., Zhou, X., & Chen, J. F. (2024). Targeting adenosine A(2A) receptors for early intervention of retinopathy of prematurity [Review]. Purinergic Signal. 10.1007/s11302-024-09986-x

D’Souza, S., & Lang, R. A. (2020). Retinal ganglion cell interactions shape the developing mammalian visual system [Review]. Development, 147(23). 10.1242/dev.196535

D’Souza, S. P., Upton, B. A., Eldred, K. C., Glass, I., Nayak, G., Grover, K., Ahmed, A., Nguyen, M. T., Hu, Y. C., Gamlin, P., & Lang, R. A. (2024). Developmental control of rod number via a light-dependent retrograde pathway from intrinsically photosensitive retinal ganglion cells. Dev Cell, 59(21), 2897–2911 e2896. 10.1016/j.devcel.2024.07.018

Duan, L.-J., Jiang, Y., & Fong, G.-H. (2024). Endothelial HIF2α suppresses retinal angiogenesis in neonatal mice by upregulating NOTCH signaling. Development, 151(11). 10.1242/dev.202802

Duan, L. J., Jiang, Y., Shi, Y., & Fong, G. H. (2023). Tailless and hypoxia inducible factor-2alpha cooperate to sustain proangiogenic states of retinal astrocytes in neonatal mice. Biol Open, 12(1). 10.1242/bio.059684

Duan, L. J., Pan, S. J., Sato, T. N., & Fong, G. H. (2017). Retinal Angiogenesis Regulates Astrocytic Differentiation in Neonatal Mouse Retinas by Oxygen Dependent Mechanisms. Sci Rep, 7(1), 17608. 10.1038/s41598-017-17962-2

Duan, L. J., Takeda, K., & Fong, G. H. (2014). Hypoxia inducible factor-2alpha regulates the development of retinal astrocytic network by maintaining adequate supply of astrocyte progenitors. PLoS One, 9(1), e84736. 10.1371/journal.pone.0084736

Duda, S., Block, C. T., Pradhan, D. R., Arzhangnia, Y., Klaiber, A., Greschner, M., & Puller, C. (2025). Spatial distribution and functional integration of displaced retinal ganglion cells. Sci Rep, 15(1), 7123. 10.1038/s41598-025-91045-5

Garcia-Gil, M., Camici, M., Allegrini, S., Pesi, R., & Tozzi, M. G. (2021). Metabolic Aspects of Adenosine Functions in the Brain. Front Pharmacol, 12, 672182. 10.3389/fphar.2021.672182

Hao, W., Wenzel, A., Obin, M. S., Chen, C. K., Brill, E., Krasnoperova, N. V., Eversole-Cire, P., Kleyner, Y., Taylor, A., Simon, M. I., Grimm, C., Reme, C. E., & Lem, J. (2002). Evidence for two apoptotic pathways in light-induced retinal degeneration. Nat Genet, 32(2), 254–260. 10.1038/ng984

He, J., Yamamoto, M., Sumiyama, K., Konagaya, Y., Terai, K., Matsuda, M., & Sato, S. (2021). Two-photon AMPK and ATP imaging reveals the bias between rods and cones in glycolysis utility. FASEB J, 35(9), e21880. 10.1096/fj.202101121R

Huang, P. C., Hsiao, Y. T., Kao, S. Y., Chen, C. F., Chen, Y. C., Chiang, C. W., Lee, C. F., Lu, J. C., Chern, Y., & Wang, C. T. (2014). Adenosine A(2A) receptor up-regulates retinal wave frequency via starburst amacrine cells in the developing rat retina. PLoS One, 9(4), e95090. 10.1371/journal.pone.0095090

Jiang, Y., Duan, L.-J., & Fong, G.-H. (2021). Oxygen-sensing mechanisms in development and tissue repair [Review]. Development, 148(23). 10.1242/dev.200030

Joyal, J. S., Gantner, M. L., & Smith, L. E. H. (2018). Retinal energy demands control vascular supply of the retina in development and disease: The role of neuronal lipid and glucose metabolism [Review]. Prog Retin Eye Res, 64, 131–156. 10.1016/j.preteyeres.2017.11.002

Koso, H., Minami, C., Tabata, Y., Inoue, M., Sasaki, E., Satoh, S., & Watanabe, S. (2009). CD73, a novel cell surface antigen that characterizes retinal photoreceptor precursor cells. Invest Ophthalmol Vis Sci, 50(11), 5411–5418. 10.1167/iovs.08-3246

Kurihara, T., Westenskow, P. D., Gantner, M. L., Usui, Y., Schultz, A., Bravo, S., Aguilar, E., Wittgrove, C., Friedlander, M., Paris, L. P., Chew, E., Siuzdak, G., & Friedlander, M. (2016). Hypoxia-induced metabolic stress in retinal pigment epithelial cells is sufficient to induce photoreceptor degeneration. Elife, 5. 10.7554/eLife.14319

Kvanta, A., Seregard, S., Sejersen, S., Kull, B., & Fredholm, B. B. (1997). Localization of adenosine receptor messenger RNAs in the rat eye. Exp Eye Res, 65(5), 595–602. 10.1006/exer.1996.0352

Lakowski, J., Han, Y. T., Pearson, R. A., Gonzalez-Cordero, A., West, E. L., Gualdoni, S., Barber, A. C., Hubank, M., Ali, R. R., & Sowden, J. C. (2011). Effective transplantation of photoreceptor precursor cells selected via cell surface antigen expression. Stem Cells, 29(9), 1391–1404. 10.1002/stem.694

Lazarus, M., Oishi, Y., Bjorness, T. E., & Greene, R. W. (2019). Gating and the Need for Sleep: Dissociable Effects of Adenosine A(1) and A(2A) Receptors [Review]. Front Neurosci, 13, 740. 10.3389/fnins.2019.00740

Li, H., Zhang, Z., Blackburn, M. R., Wang, S. W., Ribelayga, C. P., & O’Brien, J. (2013). Adenosine and dopamine receptors coregulate photoreceptor coupling via gap junction phosphorylation in mouse retina. J Neurosci, 33(7), 3135–3150. 10.1523/JNEUROSCI.2807-12.2013

Liang, J. H., Akhanov, V., Ho, A., Tawfik, M., D’Souza, S. P., Cameron, M. A., Lang, R. A., & Samuel, M. A. (2023). Dopamine signaling from ganglion cells directs layer-specific angiogenesis in the retina. Curr Biol, 33(18), 3821–3834 e3825. 10.1016/j.cub.2023.07.040

Liu, X. L., Zhou, R., Pan, Q. Q., Jia, X. L., Gao, W. N., Wu, J., Lin, J., & Chen, J. F. (2010). Genetic inactivation of the adenosine A2A receptor attenuates pathologic but not developmental angiogenesis in the mouse retina. Invest Ophthalmol Vis Sci, 51(12), 6625–6632. 10.1167/iovs.09-4900

Liu, Z., Yan, S., Wang, J., Xu, Y., Wang, Y., Zhang, S., Xu, X., Yang, Q., Zeng, X., Zhou, Y., Gu, X., Lu, S., Fu, Z., Fulton, D. J., Weintraub, N. L., Caldwell, R. B., Zhang, W., Wu, C., Liu, X. L.,…Huo, Y. (2017). Endothelial adenosine A2a receptor-mediated glycolysis is essential for pathological retinal angiogenesis. Nat Commun, 8(1), 584. 10.1038/s41467-017-00551-2

Losenkova, K., Takeda, A., Ragauskas, S., Cerrada-Gimenez, M., Vahatupa, M., Kaja, S., Paul, M. L., Schmies, C. C., Rolshoven, G., Muller, C. E., Sandholm, J., Jalkanen, S., Kalesnykas, G., & Yegutkin, G. G. (2022). CD73 controls ocular adenosine levels and protects retina from light-induced phototoxicity. Cell Mol Life Sci, 79(3), 152. 10.1007/s00018-022-04187-4

Mehalow, A. K., Kameya, S., Smith, R. S., Hawes, N. L., Denegre, J. M., Young, J. A., Bechtold, L., Haider, N. B., Tepass, U., Heckenlively, J. R., Chang, B., Naggert, J. K., & Nishina, P. M. (2003). CRB1 is essential for external limiting membrane integrity and photoreceptor morphogenesis in the mammalian retina. Hum Mol Genet, 12(17), 2179–2189. 10.1093/hmg/ddg232

Miwa, Y., Tsubota, K., & Kurihara, T. (2019). Effect of midazolam, medetomidine, and butorphanol tartrate combination anesthetic on electroretinograms of mice. Molecular vision, 25, 645–653. http://www.molvis.org/molvis/v25/645

Nakamura-Ishizu, A., Kurihara, T., Okuno, Y., Ozawa, Y., Kishi, K., Goda, N., Tsubota, K., Okano, H., Suda, T., & Kubota, Y. (2012). The formation of an angiogenic astrocyte template is regulated by the neuroretina in a HIF-1-dependent manner. Dev Biol, 363(1), 106–114. 10.1016/j.ydbio.2011.12.027

Nguyen, M. T., Vemaraju, S., Nayak, G., Odaka, Y., Buhr, E. D., Alonzo, N., Tran, U., Batie, M., Upton, B. A., Darvas, M., Kozmik, Z., Rao, S., Hegde, R. S., Iuvone, P. M., Van Gelder, R. N., & Lang, R. A. (2019). An opsin 5-dopamine pathway mediates light-dependent vascular development in the eye. Nat Cell Biol, 21(4), 420–429. 10.1038/s41556-019-0301-x

Oishi, Y., Xu, Q., Wang, L., Zhang, B. J., Takahashi, K., Takata, Y., Luo, Y. J., Cherasse, Y., Schiffmann, S. N., de Kerchove d’Exaerde, A., Urade, Y., Qu, W. M., Huang, Z. L., & Lazarus, M. (2017). Slow-wave sleep is controlled by a subset of nucleus accumbens core neurons in mice. Nat Commun, 8(1), 734. 10.1038/s41467-017-00781-4

Organisciak, D. T., & Vaughan, D. K. (2010). Retinal light damage: mechanisms and protection [Review]. Prog Retin Eye Res, 29(2), 113–134. 10.1016/j.preteyeres.2009.11.004

Paisley, C. E., & Kay, J. N. (2021). Seeing stars: Development and function of retinal astrocytes [Review]. Dev Biol, 478, 144–154. 10.1016/j.ydbio.2021.07.007

Pearson, R. A., Dale, N., Llaudet, E., & Mobbs, P. (2005). ATP released via gap junction hemichannels from the pigment epithelium regulates neural retinal progenitor proliferation. Neuron, 46(5), 731–744. 10.1016/j.neuron.2005.04.024

Pugh, E. N., Jr., & Lamb, T. D. (1993). Amplification and kinetics of the activation steps in phototransduction. Biochim Biophys Acta, 1141(2-3), 111–149. 10.1016/0005-2728(93)90038-h

Rao, S., Chun, C., Fan, J., Kofron, J. M., Yang, M. B., Hegde, R. S., Ferrara, N., Copenhagen, D. R., & Lang, R. A. (2013). A direct and melanopsin-dependent fetal light response regulates mouse eye development. Nature, 494(7436), 243–246. 10.1038/nature11823

Ribelayga, C., & Mangel, S. C. (2005). A circadian clock and light/dark adaptation differentially regulate adenosine in the mammalian retina. J Neurosci, 25(1), 215–222. 10.1523/JNEUROSCI.3138-04.2005

Sakurai, K., Onishi, A., Imai, H., Chisaka, O., Ueda, Y., Usukura, J., Nakatani, K., & Shichida, Y. (2007). Physiological properties of rod photoreceptor cells in green-sensitive cone pigment knock-in mice. J Gen Physiol, 130(1), 21–40. 10.1085/jgp.200609729

Santiago, A. R., Madeira, M. H., Boia, R., Aires, I. D., Rodrigues-Neves, A. C., Santos, P. F., & Ambrosio, A. F. (2020). Keep an eye on adenosine: Its role in retinal inflammation [Review]. Pharmacol Ther, 210, 107513. 10.1016/j.pharmthera.2020.107513

Sarin, S., Zuniga-Sanchez, E., Kurmangaliyev, Y. Z., Cousins, H., Patel, M., Hernandez, J., Zhang, K. X., Samuel, M. A., Morey, M., Sanes, J. R., & Zipursky, S. L. (2018). Role for Wnt Signaling in Retinal Neuropil Development: Analysis via RNA-Seq and In Vivo Somatic CRISPR Mutagenesis. Neuron, 98(1), 109–126 e108. 10.1016/j.neuron.2018.03.004

Sato, Y., & Yanagita, M. (2013). Renal anemia: from incurable to curable. Am J Physiol Renal Physiol, 305(9), F1239–1248. 10.1152/ajprenal.00233.2013

Semba, H., Takeda, N., Isagawa, T., Sugiura, Y., Honda, K., Wake, M., Miyazawa, H., Yamaguchi, Y., Miura, M., Jenkins, D. M., Choi, H., Kim, J. W., Asagiri, M., Cowburn, A. S., Abe, H., Soma, K., Koyama, K., Katoh, M., Sayama, K.,…Komuro, I. (2016). HIF-1alpha-PDK1 axis-induced active glycolysis plays an essential role in macrophage migratory capacity. Nat Commun, 7, 11635. 10.1038/ncomms11635

Stella, S. L., Jr., Bryson, E. J., Cadetti, L., & Thoreson, W. B. (2003). Endogenous adenosine reduces glutamatergic output from rods through activation of A2-like adenosine receptors. J Neurophysiol, 90(1), 165–174. 10.1152/jn.00671.2002

Stone, J., Maslim, J., Valter-Kocsi, K., Mervin, K., Bowers, F., Chu, Y., Barnett, N., Provis, J., Lewis, G., Fisher, S. K., Bisti, S., Gargini, C., Cervetto, L., Merin, S., & Peer, J. (1999). Mechanisms of photoreceptor death and survival in mammalian retina [Review]. Prog Retin Eye Res, 18(6), 689–735. 10.1016/s1350-9462(98)00032-9

Swaroop, A., Kim, D., & Forrest, D. (2010). Transcriptional regulation of photoreceptor development and homeostasis in the mammalian retina [Review]. Nat Rev Neurosci, 11(8), 563–576. 10.1038/nrn2880

Syed, M. M., Lee, S., Zheng, J., & Zhou, Z. J. (2004). Stage-dependent dynamics and modulation of spontaneous waves in the developing rabbit retina. J Physiol, 560(Pt 2), 533–549. 10.1113/jphysiol.2004.066597

Synnestvedt, K., Furuta, G. T., Comerford, K. M., Louis, N., Karhausen, J., Eltzschig, H. K., Hansen, K. R., Thompson, L. F., & Colgan, S. P. (2002). Ecto-5’-nucleotidase (CD73) regulation by hypoxia-inducible factor-1 mediates permeability changes in intestinal epithelia. J Clin Invest, 110(7), 993–1002. 10.1172/JCI15337

Tao, C., & Zhang, X. (2014). Development of astrocytes in the vertebrate eye. Dev Dyn, 243(12), 1501–1510. 10.1002/dvdy.24190

Tiriac, A., Smith, B. E., & Feller, M. B. (2018). Light Prior to Eye Opening Promotes Retinal Waves and Eye-Specific Segregation. Neuron, 100(5), 1059–1065 e1054. 10.1016/j.neuron.2018.10.011

Torborg, C. L., & Feller, M. B. (2005). Spontaneous patterned retinal activity and the refinement of retinal projections [Review]. Prog Neurobiol, 76(4), 213–235. 10.1016/j.pneurobio.2005.09.002

van de Pavert, S. A., Kantardzhieva, A., Malysheva, A., Meuleman, J., Versteeg, I., Levelt, C., Klooster, J., Geiger, S., Seeliger, M. W., Rashbass, P., Le Bivic, A., & Wijnholds, J. (2004). Crumbs homologue 1 is required for maintenance of photoreceptor cell polarization and adhesion during light exposure. J Cell Sci, 117(Pt 18), 4169–4177. 10.1242/jcs.01301

van de Pavert, S. A., Sanz, A. S., Aartsen, W. M., Vos, R. M., Versteeg, I., Beck, S. C., Klooster, J., Seeliger, M. W., & Wijnholds, J. (2007). Crb1 is a determinant of retinal apical Muller glia cell features. Glia, 55(14), 1486–1497. 10.1002/glia.20561

Wang, S., Sengel, C., Emerson, M. M., & Cepko, C. L. (2014). A gene regulatory network controls the binary fate decision of rod and bipolar cells in the vertebrate retina. Dev Cell, 30(5), 513–527. 10.1016/j.devcel.2014.07.018

Wei, C. J., Li, W., & Chen, J. F. (2011). Normal and abnormal functions of adenosine receptors in the central nervous system revealed by genetic knockout studies [Review]. Biochim Biophys Acta, 1808(5), 1358–1379. 10.1016/j.bbamem.2010.12.018

Weidemann, A., Krohne, T. U., Aguilar, E., Kurihara, T., Takeda, N., Dorrell, M. I., Simon, M. C., Haase, V. H., Friedlander, M., & Johnson, R. S. (2010). Astrocyte hypoxic response is essential for pathological but not developmental angiogenesis of the retina. Glia, 58(10), 1177–1185. 10.1002/glia.20997

Weiner, G. A., Shah, S. H., Angelopoulos, C. M., Bartakova, A. B., Pulido, R. S., Murphy, A., Nudleman, E., Daneman, R., & Goldberg, J. L. (2019). Cholinergic neural activity directs retinal layer-specific angiogenesis and blood retinal barrier formation. Nat Commun, 10(1), 2477. 10.1038/s41467-019-10219-8

Wenzel, A., Grimm, C., Samardzija, M., & Reme, C. E. (2005). Molecular mechanisms of light-induced photoreceptor apoptosis and neuroprotection for retinal degeneration [Review]. Prog Retin Eye Res, 24(2), 275–306. 10.1016/j.preteyeres.2004.08.002

West, H., Richardson, W. D., & Fruttiger, M. (2005). Stabilization of the retinal vascular network by reciprocal feedback between blood vessels and astrocytes. Development, 132(8), 1855–1862. 10.1242/dev.01732

Wu, Z., Cui, Y., Wang, H., Wu, H., Wan, Y., Li, B., Wang, L., Pan, S., Peng, W., Dong, A., Yuan, Z., Jing, M., Xu, M., Luo, M., & Li, Y. (2023). Neuronal activity-induced, equilibrative nucleoside transporter-dependent, somatodendritic adenosine release revealed by a GRAB sensor. Proc Natl Acad Sci U S A, 120(14), e2212387120. 10.1073/pnas.2212387120

Wurm, A., Lipp, S., Pannicke, T., Linnertz, R., Krugel, U., Schulz, A., Farber, K., Zahn, D., Grosse, J., Wiedemann, P., Chen, J., Schoneberg, T., Illes, P., Reichenbach, A., & Bringmann, A. (2010). Endogenous purinergic signaling is required for osmotic volume regulation of retinal glial cells. J Neurochem, 112(5), 1261–1272. 10.1111/j.1471-4159.2009.06541.x

Wurm, A., Pannicke, T., Iandiev, I., Francke, M., Hollborn, M., Wiedemann, P., Reichenbach, A., Osborne, N. N., & Bringmann, A. (2011). Purinergic signaling involved in Muller cell function in the mammalian retina [Review]. Prog Retin Eye Res, 30(5), 324–342. 10.1016/j.preteyeres.2011.06.001

Xu, Y., Wang, Y., Yan, S., Zhou, Y., Yang, Q., Pan, Y., Zeng, X., An, X., Liu, Z., Wang, L., Xu, J., Cao, Y., Fulton, D. J., Weintraub, N. L., Bagi, Z., Hoda, M. N., Wang, X., Li, Q., Hong, M.,…Huo, Y. (2017). Intracellular adenosine regulates epigenetic programming in endothelial cells to promote angiogenesis. EMBO Mol Med, 9(9), 1263–1278. 10.15252/emmm.201607066

Yan, W., Laboulaye, M. A., Tran, N. M., Whitney, I. E., Benhar, I., & Sanes, J. R. (2020). Mouse Retinal Cell Atlas: Molecular Identification of over Sixty Amacrine Cell Types. J Neurosci, 40(27), 5177–5195. 10.1523/JNEUROSCI.0471-20.2020

Yuan, X., Ruan, W., Bobrow, B., Carmeliet, P., & Eltzschig, H. K. (2024). Targeting hypoxia-inducible factors: therapeutic opportunities and challenges [Review]. Nat Rev Drug Discov, 23(3), 175–200. 10.1038/s41573-023-00848-6

Zhang, J., Roberts, J. M., Chang, F., Schwakopf, J., & Vetter, M. L. (2023). Jarid2 promotes temporal progression of retinal progenitors via repression of Foxp1. Cell Rep, 42(3), 112237. 10.1016/j.celrep.2023.112237

Zhang, S., Li, B., Tang, L., Tong, M., Jiang, N., Gu, X., Zhang, Y., Ge, Y., Liu, X. L., & Chen, J. F. (2022). Disruption of CD73-Derived and Equilibrative sNucleoside Transporter 1-Mediated Adenosine Signaling Exacerbates Oxygen-Induced Retinopathy. Am J Pathol. 10.1016/j.ajpath.2022.07.014

Zhang, S., Li, H., Li, B., Zhong, D., Gu, X., Tang, L., Wang, Y., Wang, C., Zhou, R., Li, Y., He, Y., Chen, M., Huo, Y., Liu, X. L., & Chen, J. F. (2015). Adenosine A1 Receptors Selectively Modulate Oxygen-Induced Retinopathy at the Hyperoxic and Hypoxic Phases by Distinct Cellular Mechanisms. Invest Ophthalmol Vis Sci, 56(13), 8108–8119. 10.1167/iovs.15-17202

Zhong, D. J., Zhang, Y., Zhang, S., Ge, Y. Y., Tong, M., Feng, Y., You, F., Zhao, X., Wang, K., Zhang, L., Liu, X., & Chen, J. F. (2021). Adenosine A(2A) receptor antagonism protects against hyperoxia-induced retinal vascular loss via cellular proliferation. FASEB J, 35(9), e21842. 10.1096/fj.202100414RR

