## Supplementary figures and images for "Spatiotemporal dynamics of CD73 in mouse retina under physiological conditions"

### Figure 1-figure supplement 1

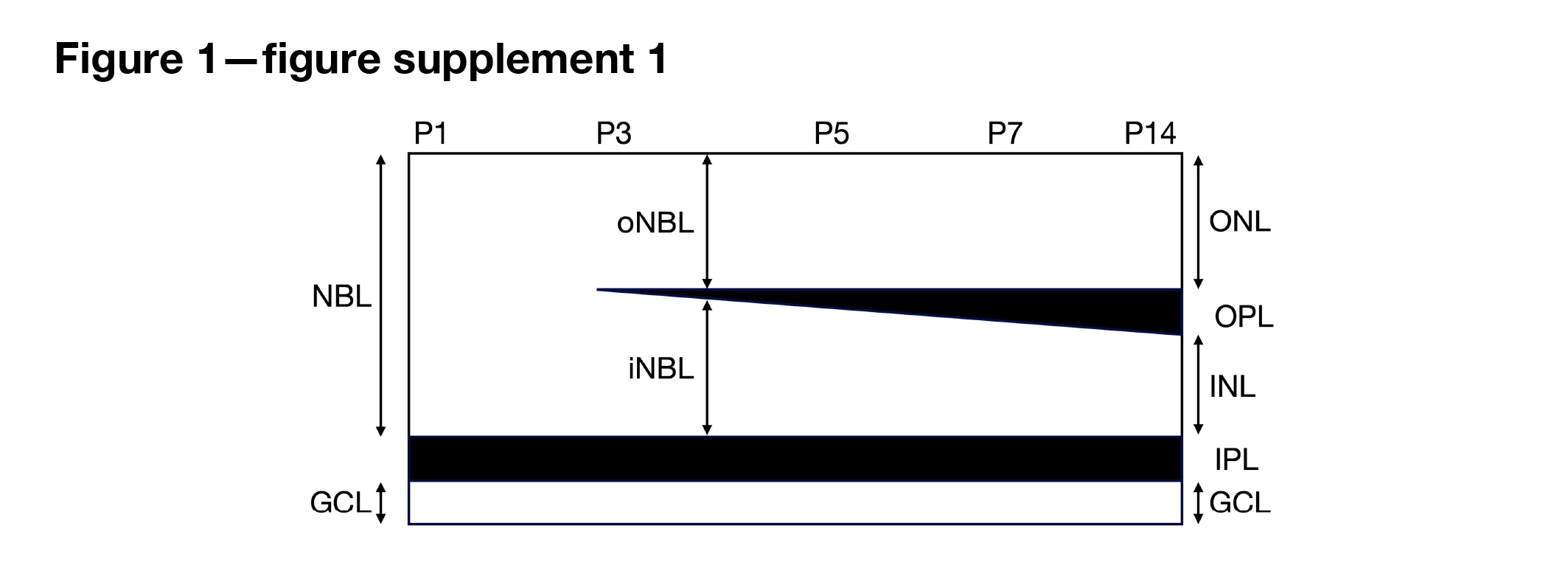

### Figure 1-figure supplement 2

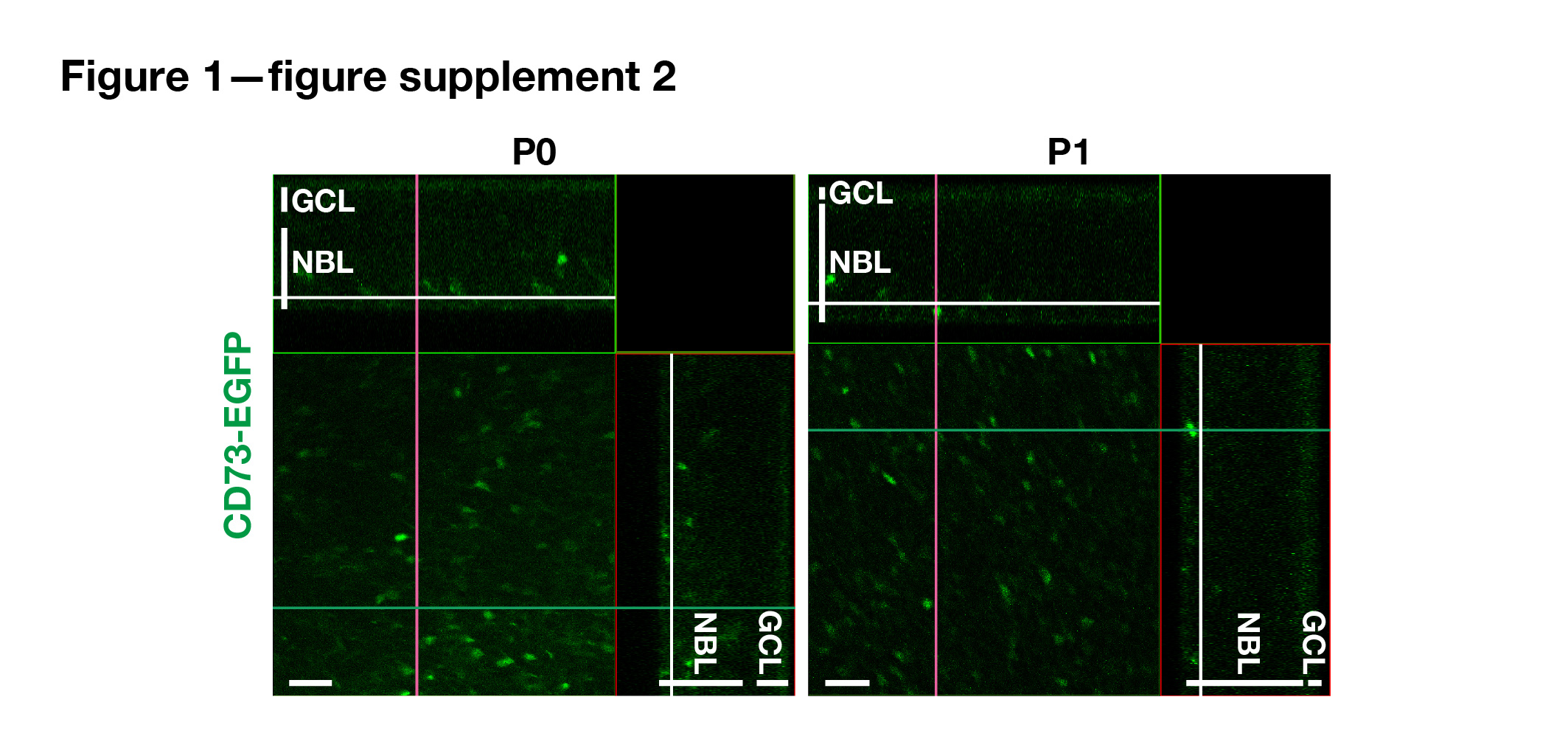

### Figure 1-figure supplement 3

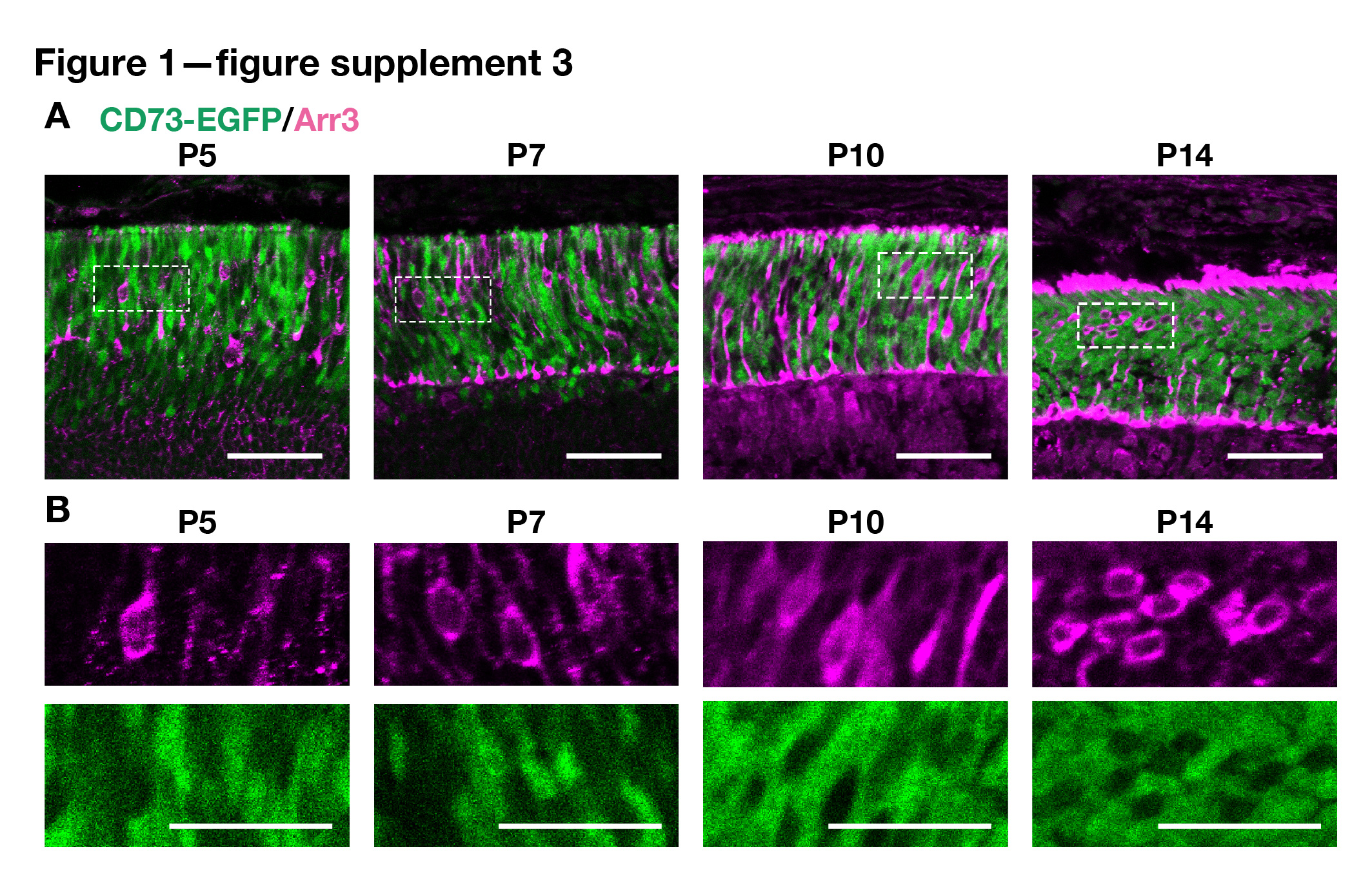

### Figure 2-figure supplement 1

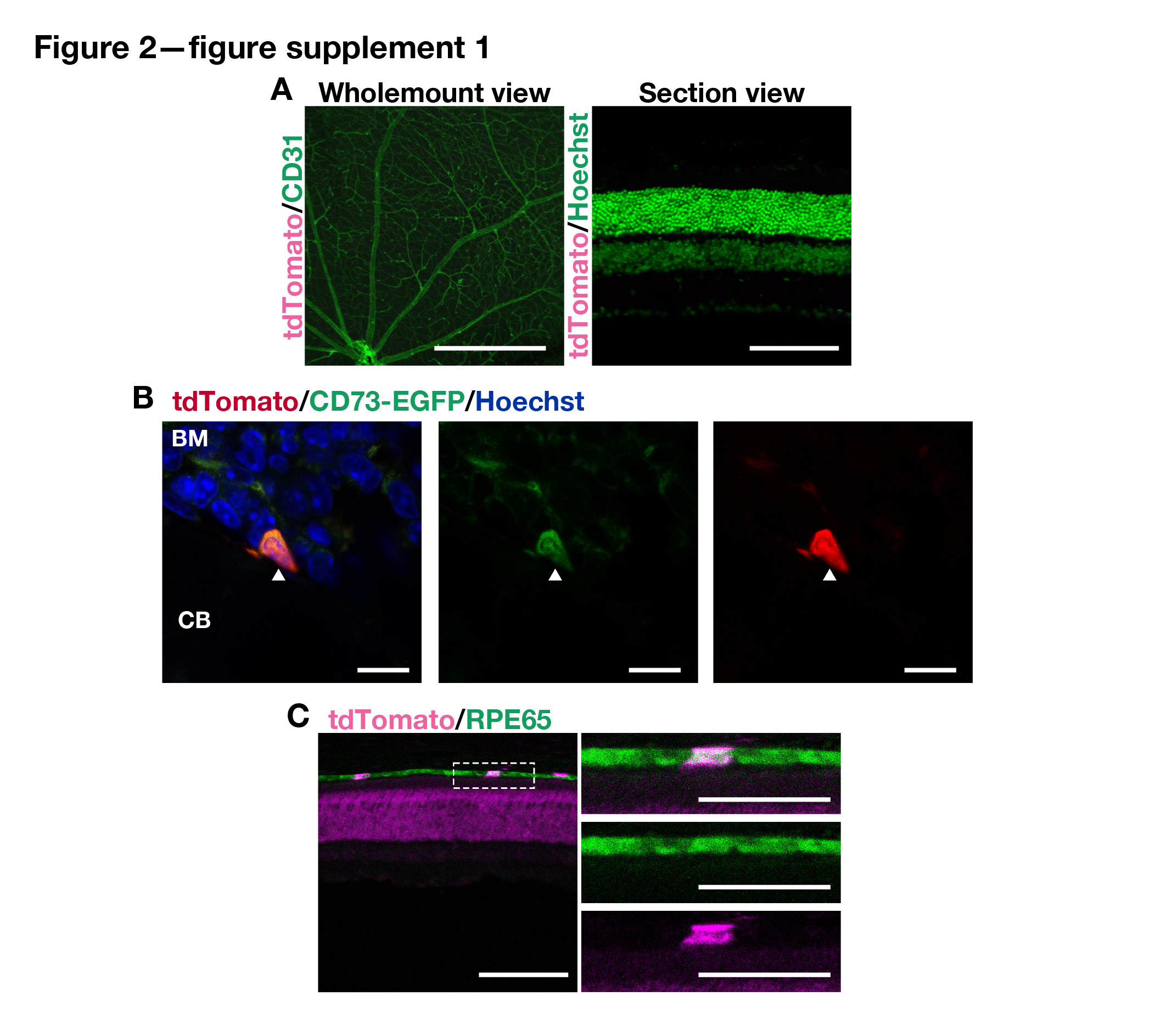

### Figure 2-figure supplement 2

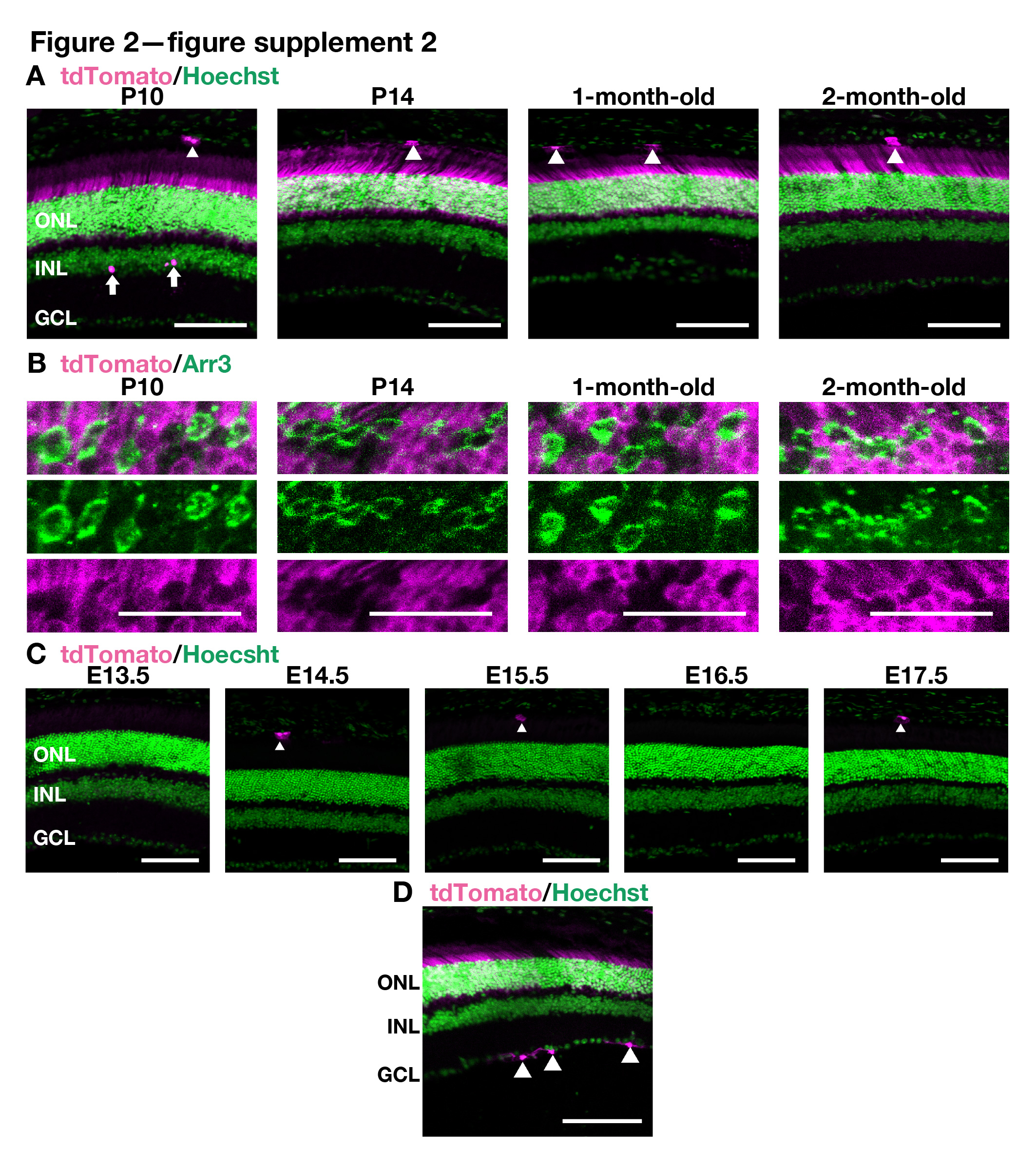

### Figure 3-figure supplement 1

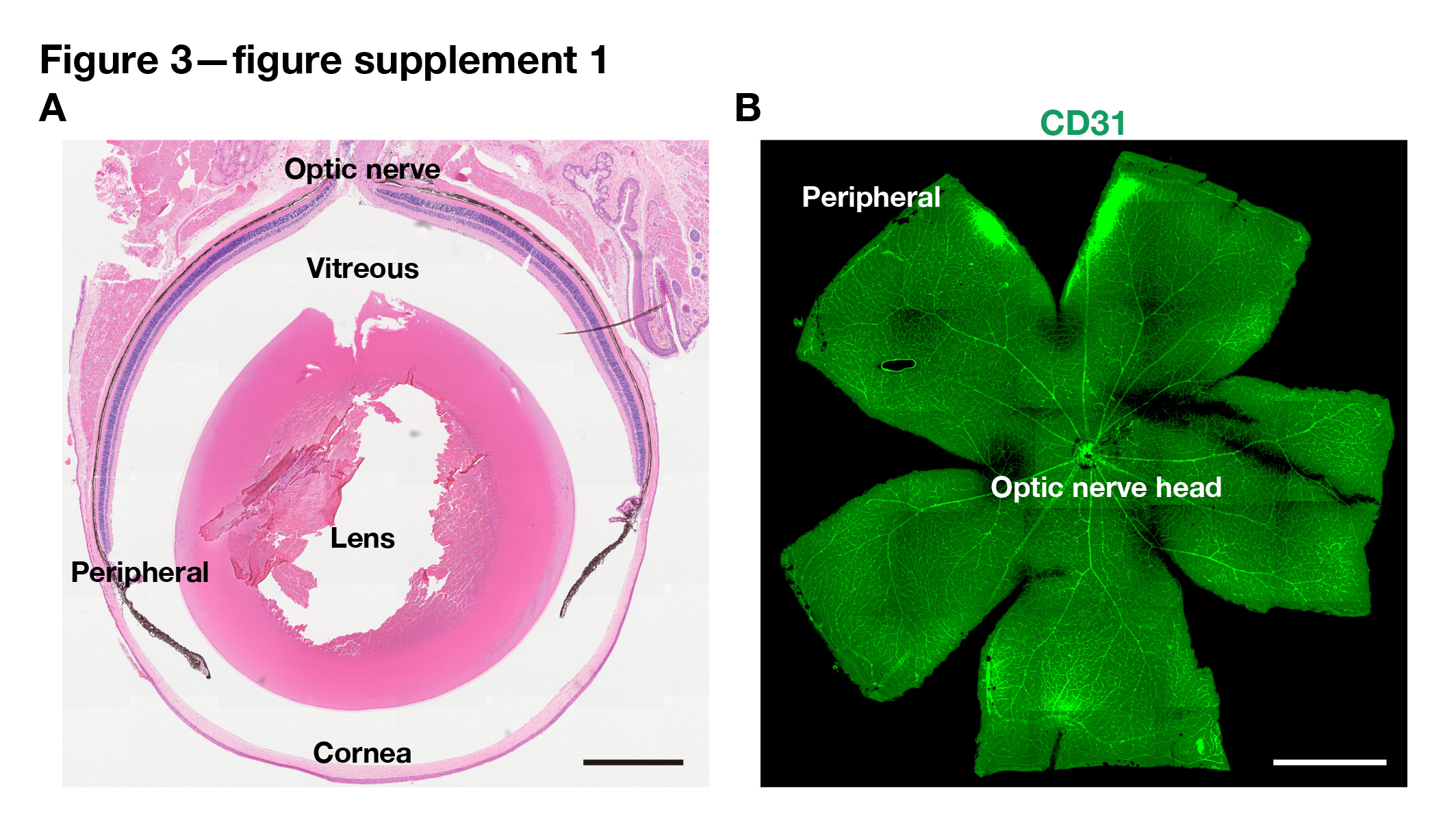

### Figure 4-figure supplement 1

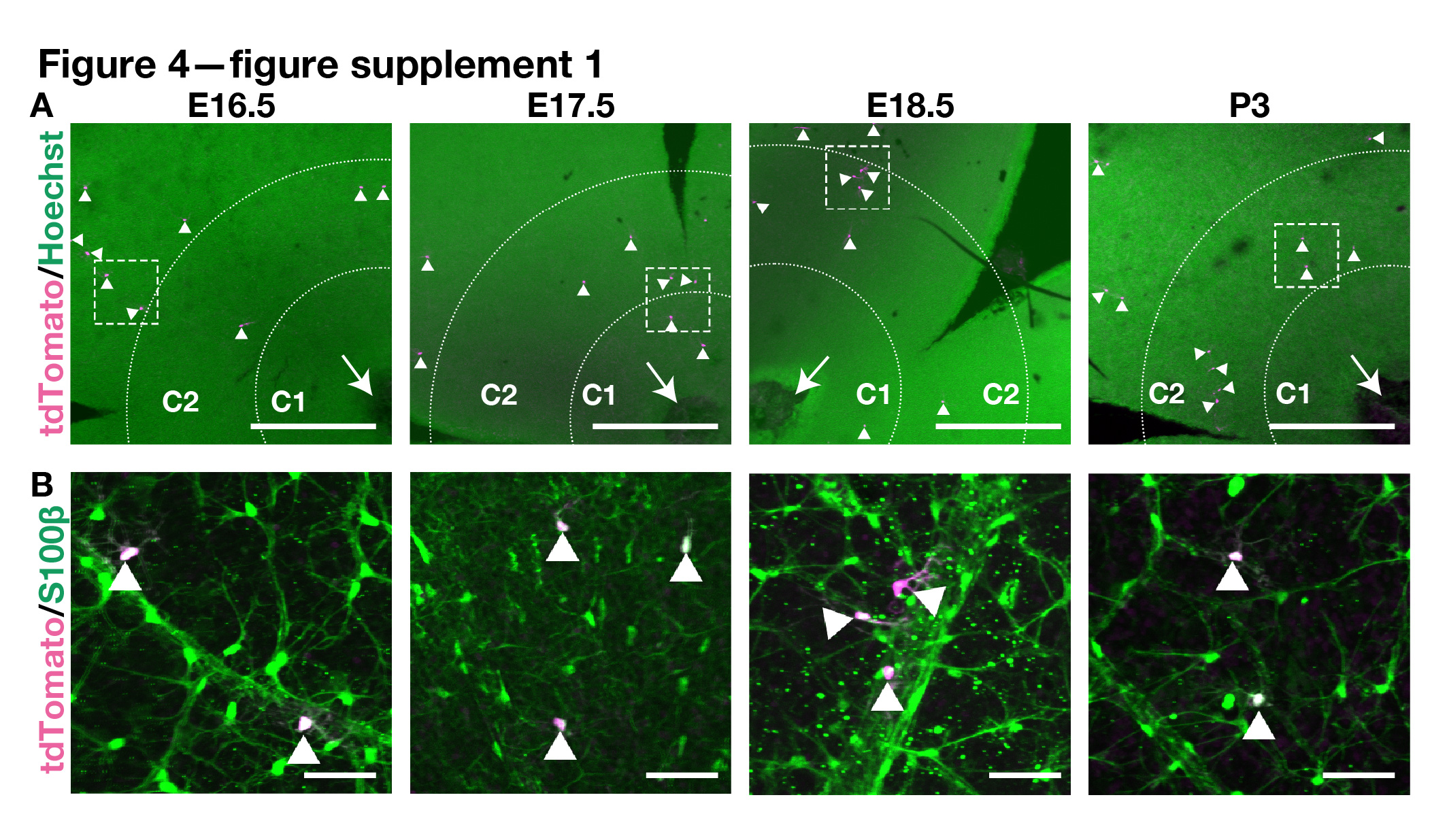

### Figure 5-figure supplement 1

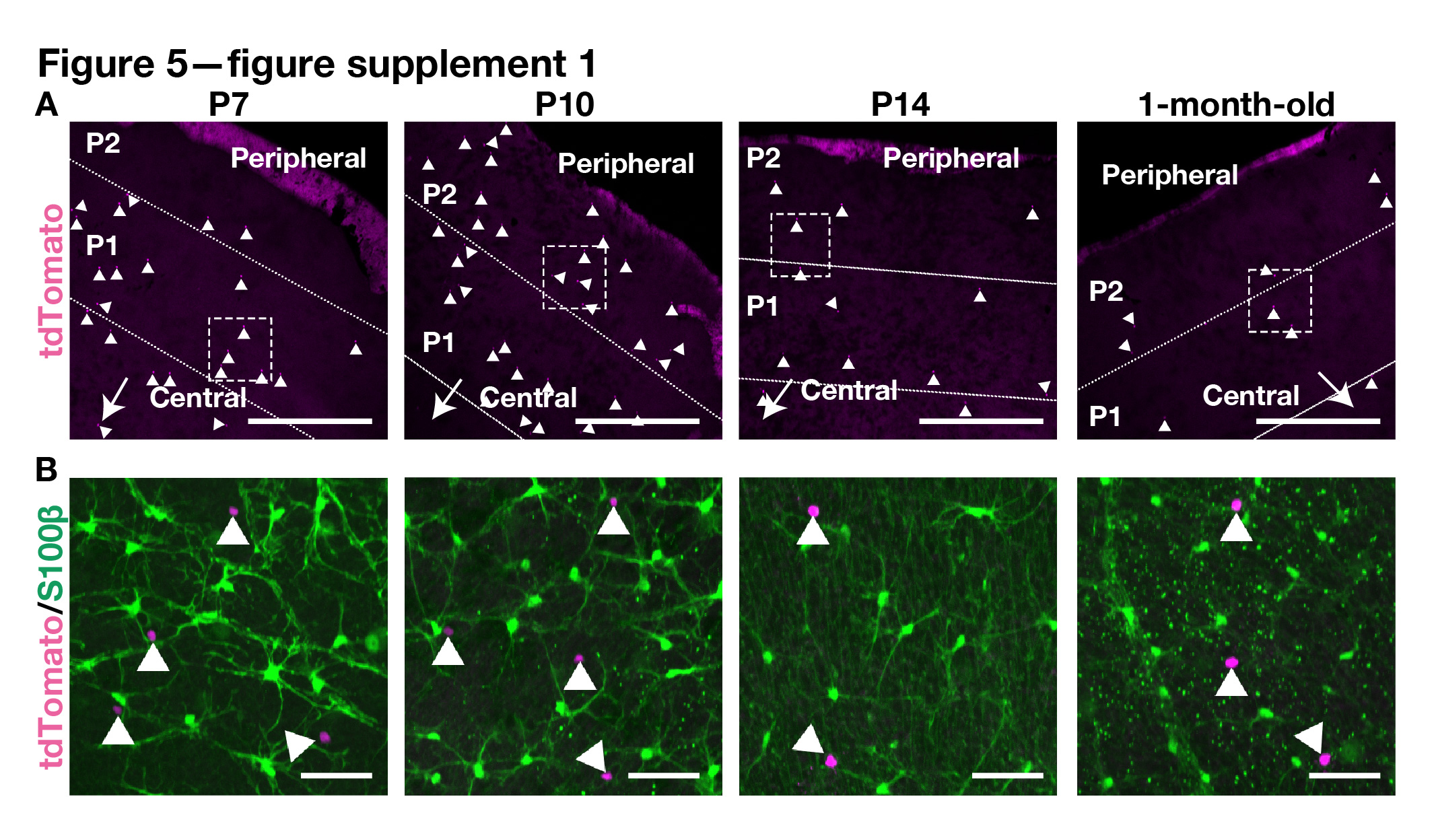

### Figure 5-figure supplement 2

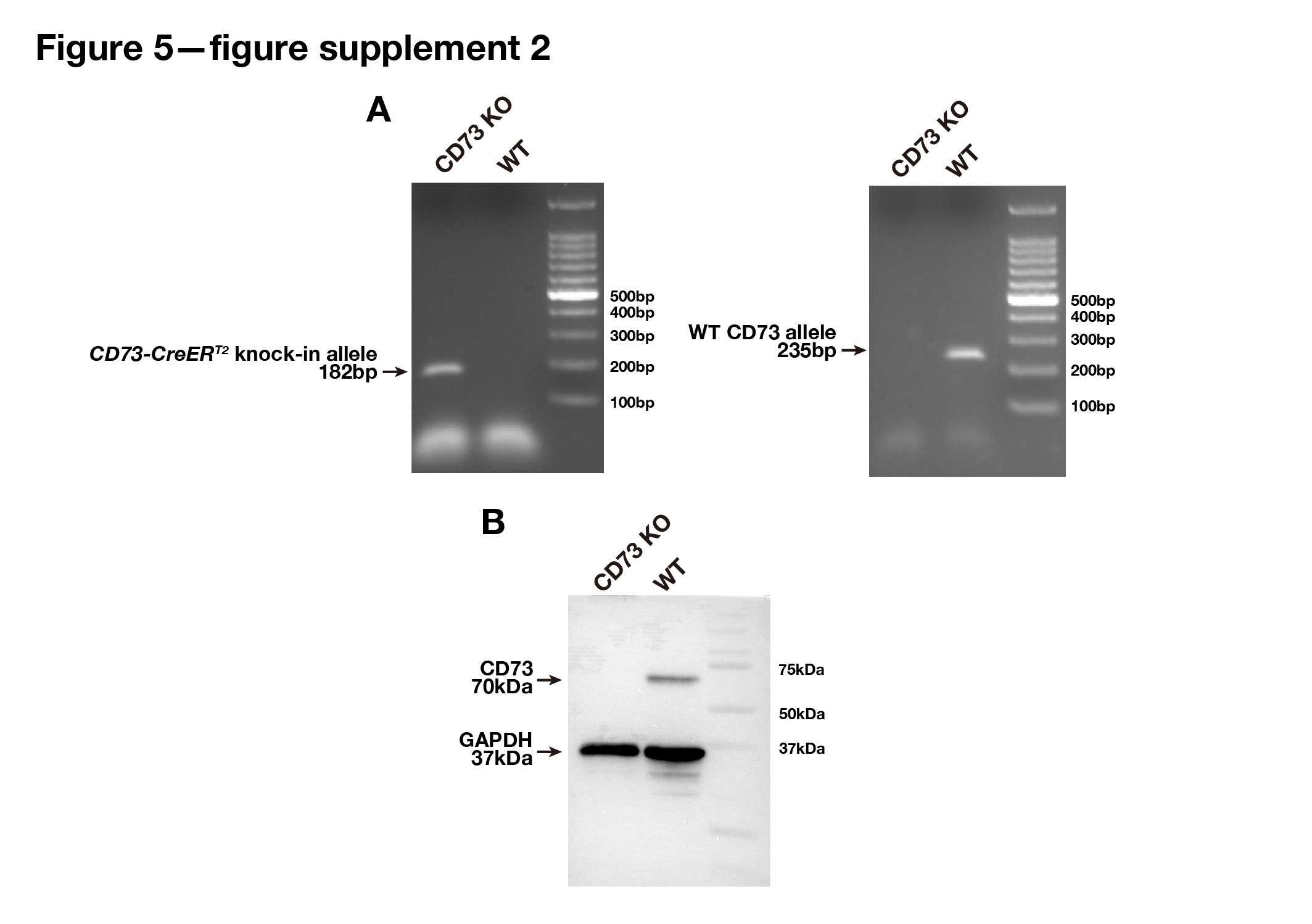

### Figure 5-figure supplement 3

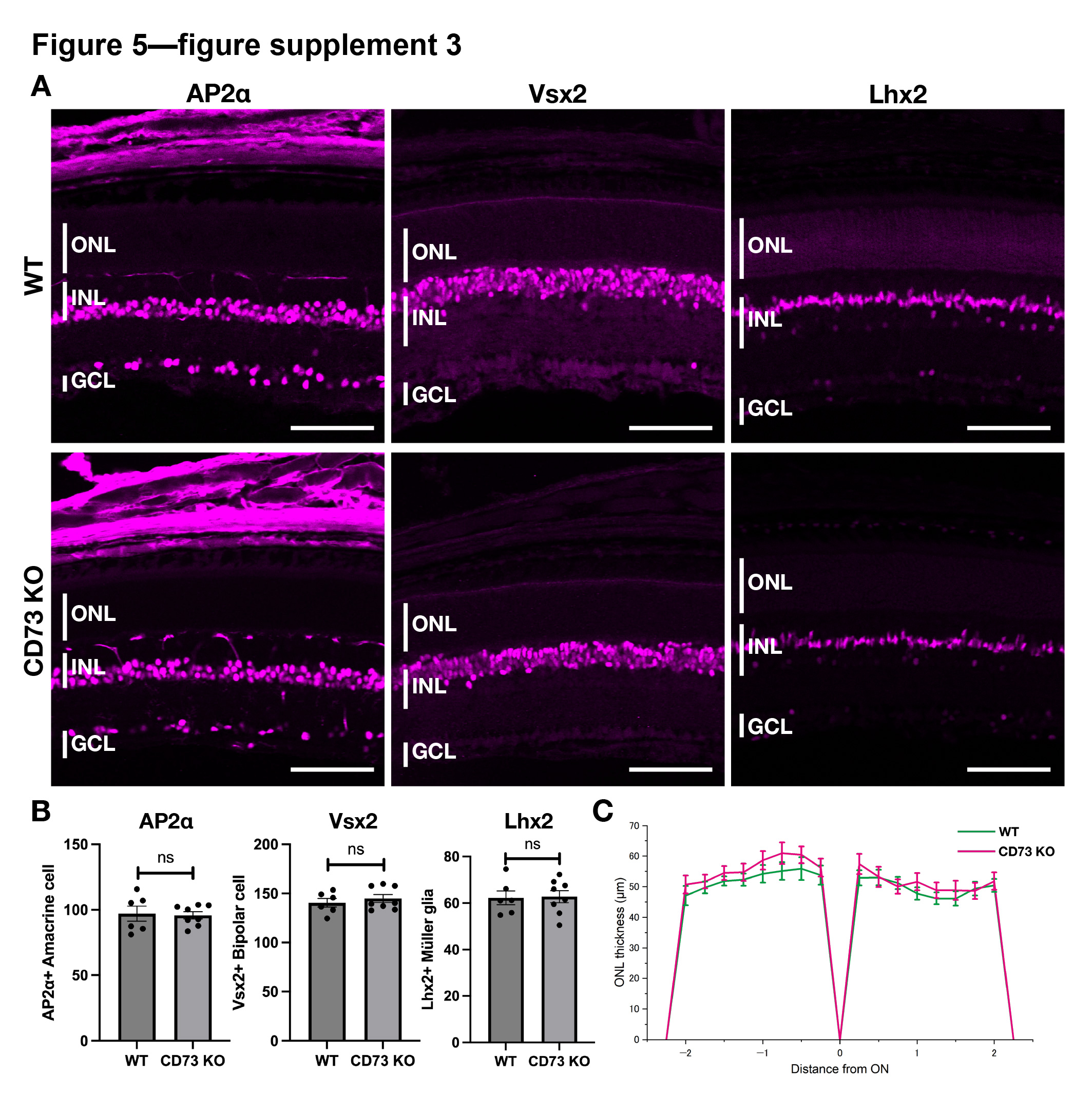

### Figure 6-figure supplement 1

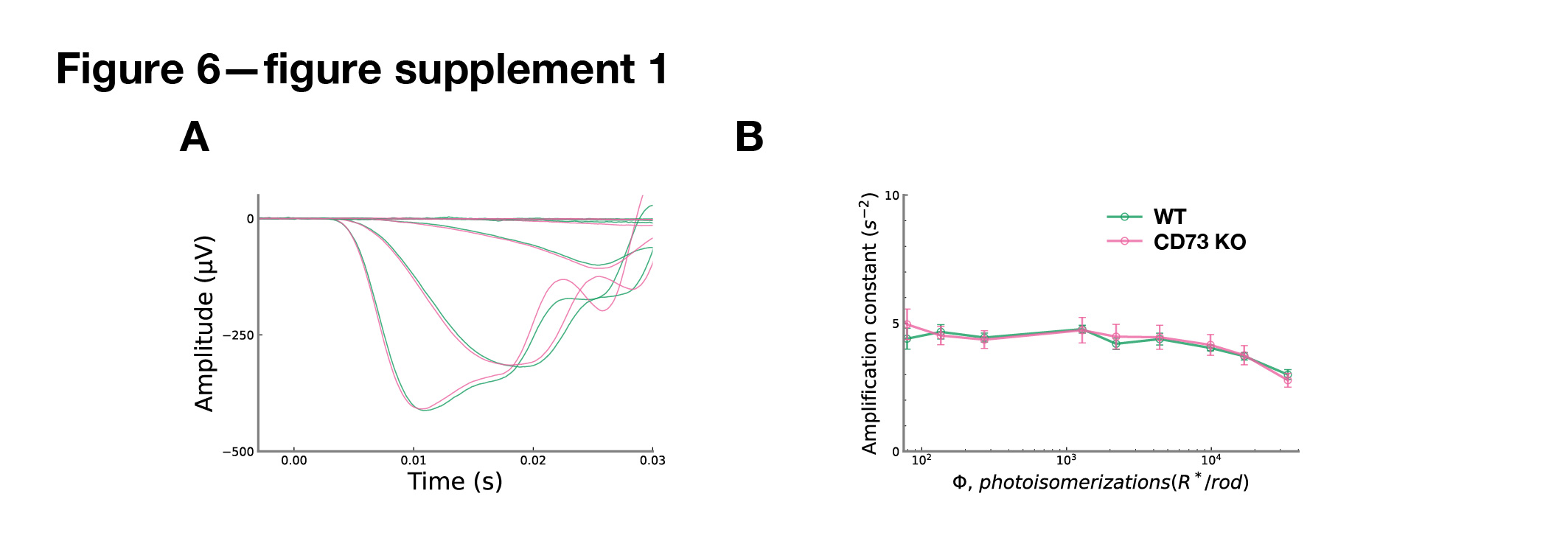

### Figure 7-figure supplement 1

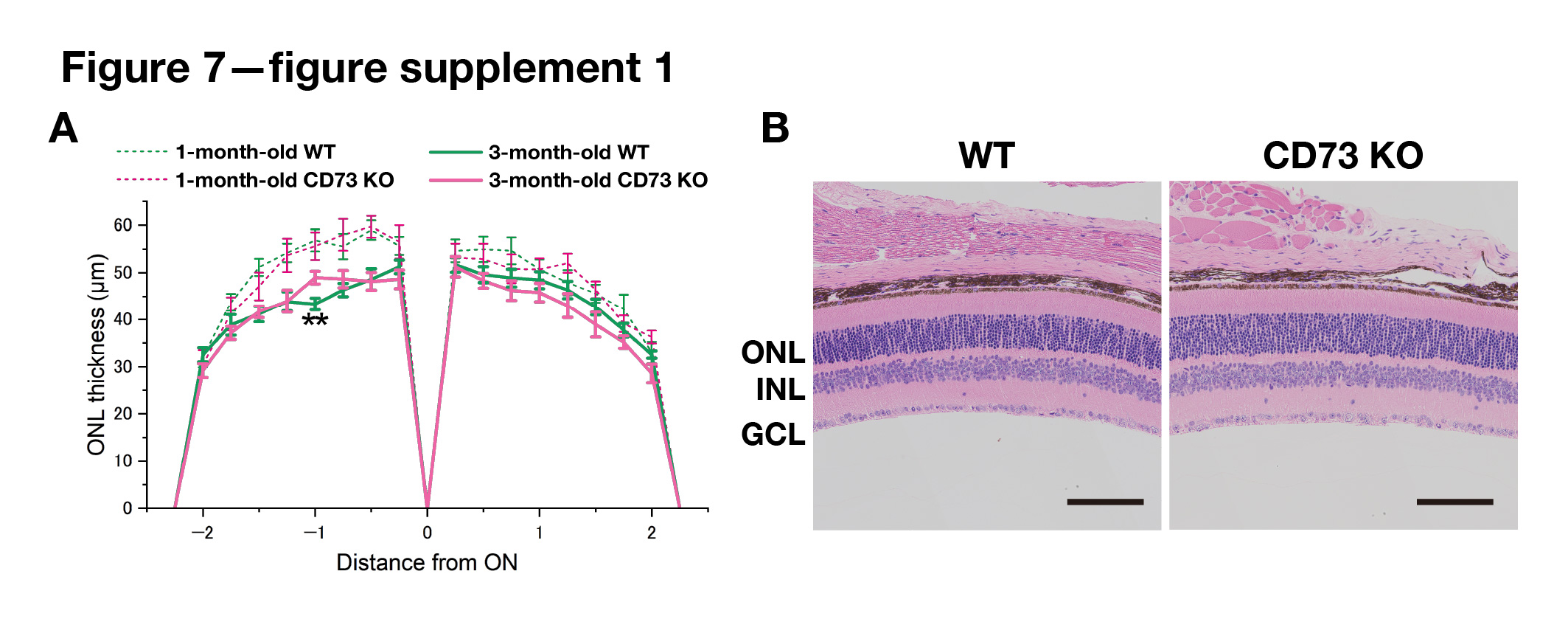
